# Climate biogeography of *Arabidopsis thaliana:* linking distribution models and individual variation

**DOI:** 10.1101/2022.03.06.483202

**Authors:** Christina Yim, Emily S. Bellis, Victoria L. DeLeo, Diana Gamba, Robert Muscarella, Jesse R. Lasky

**Author notes:** **Biosketch**. The research team is interested in understanding how individual-level processes influence evolutionary and ecological dynamics at continental scales. **Author contributions**. All authors contributed to study design. CY, ESB, VLD, DG, and JRL conducted analyses. CY and JRL led the writing with contributions from all authors.

## Abstract

**AIM:** Patterns of individual variation are key to testing hypotheses about the mechanisms underlying biogeographic patterns. However, it is challenging to gather data on individual-level variation at large spatial scales. Model organisms are potentially important systems for biogeographical studies, given the available range-wide natural history collections, and the importance of providing biogeographical context to their genetic and phenotypic diversity.

**LOCATION:** Global

**TAXON:** *Arabidopsis thaliana* (“Arabidopsis”)

**METHODS:** We fit occurrence records to climate data, and then projected the distribution of Arabidopsis under last glacial maximum, current, and future climates. We confronted model predictions with individual performance measured on 2,194 herbarium specimens, and we asked whether predicted suitability was associated with life-history and genomic variation measured on ∼900 natural accessions.

**RESULTS:** The most important climate variables constraining the Arabidopsis distribution were winter cold in northern and high elevation regions and summer heat in southern regions. Herbarium specimens from regions with lower habitat suitability in both northern and southern regions were smaller, supporting the hypothesis that the distribution of Arabidopsis is constrained by climate-associated factors. Climate anomalies partly explained interannual variation in herbarium specimen size, but these did not closely correspond to local limiting factors identified in the distribution model. Late-flowering genotypes were absent from the lowest suitability regions, suggesting slower life histories are only viable closer to the center of the realized niche. We identified glacial refugia farther north than previously recognized, as well as refugia concordant with previous population genetic findings. Lower latitude populations, known to be genetically distinct, are most threatened by future climate change. The recently colonized range of Arabidopsis was well-predicted by our native-range model applied to certain regions but not others, suggesting it has colonized novel climates.

**MAIN CONCLUSIONS:** Integration of distribution models with performance data from vast natural history collections is a route forward for testing biogeographical hypotheses about species distributions and their relationship with evolutionary fitness across large scales.

## Introduction

Biogeographic patterns emerge from processes that act on many individuals and different local populations. Individuals die or survive in response to fluctuating environments, individuals migrate, and population allele frequencies change over time. While theory has often described how processes acting on individuals and local populations can generate biogeographic patterns (Keitt et al., 2001; Kirkpatrick & Barton, 1997; Pulliam, 2000) empirical study is challenged by the logistics required to measure individual-level variation across entire species ranges.

Additionally, there is a growing call to connect range-wide, large scale model predictions with information on molecular, individual, and population-level processes (Kearney & Porter, 2009; Lasky et al., 2020). Thus there is a need for greater integration of biogeography with organismal biology to test hypotheses about the organismal and population mechanisms controlling distributions, and to understand how distributions and biogeographical history leave their imprints on individuals and populations.

In particular, a major goal in biology and biogeography is to understand how environmental conditions limit individual performance and species distributions. First, the environment-distribution relationship can help project species distributions under past climates to understand their history (Forester et al., 2013). Second, the environment-distribution relationship can identify regions with currently suitable habitat but that are unoccupied for other reasons (Elith et al., 2010). Third, the environment-distribution relationship can predict how distributions will shift under future environments (Thomas et al., 2004).

Environment-distribution relationships fundamentally arise from processes acting on individuals (Clark, 2010). On average, transplant experiments show that individual performance declines outside species’ natural range (Hargreaves et al., 2014) and efforts to integrate information at the individual level into distribution models are emerging (Buckley et al., 2011; Elith et al., 2010; Lasky et al., 2020; Merow et al., 2014; Samis & Eckert, 2007). Where populations inhabit harsh environments (e.g. at range margins), local adaptations can emerge, such as life history changes to tolerate or escape harsh periods (Bontrager et al., 2021). For example, in Arabidopsis there is evidence that local adaptation to abiotic environment involves genetic changes in life history (e.g. flowering time) (Lovell et al., 2013; Martínez-Berdeja et al., 2020).

A vast resource of individual-level information can be found in natural history collections, and advances in the digitization of museum specimens are rapidly expanding the available data on range-wide variation (Heberling, 2021; Lopez et al., 2019). For example, Bontrager & Angert (2015) showed with herbarium specimens of *Clarkia* that fecundity decreased with drier summers, and toward one range margin both summer precipitation and individual fecundity declined, suggesting a limiting mechanism. In Arabidopsis, DeLeo et al. (2020) found decadal shifts in traits of herbarium specimens. For many species, seed banks (a type of natural history collection) house great diversity from across their ranges that can also be used to study their biogeography (Estarague et al., 2021; Scholl et al., 2000). These approaches are potentially valuable in model organisms, where researchers can link detailed information on genetics and organismal biology with population and community processes (Rudman et al., 2019; Takou et al., 2019). In this spirit we focus on *Arabidopsis thaliana,* a small annual plant (hereafter referred to as “Arabidopsis”) (Koornneef & Meinke, 2010). Arabidopsis has been key to linking molecular biology, physiology, evolution, and ecology, but past biogeographic studies lacked statistical inference (Hoffmann, 2002) or overlooked large parts of its range (Banta et al., 2012; Zou et al., 2017). Distribution models fit to occurrence data are explicit quantitative statement of environment-distribution relationships and allow inference of the environmental factors limiting distributions (Elith et al., 2010; G. Li et al., 2015).

Arabidopsis is native across Eurasia and Africa and with human assistance has colonized the Americas and Australia. Other species in the genus *Arabidopsis* are more restricted to cool temperate climates, with Arabidopsis having expanded to a broader range, *e.g.* Mediterranean habitats (Hoffmann, 2005). Arabidopsis can behave as a spring annual with a rapid life cycle, germinating in the spring and flowering in the late spring and summer. However, many individuals are longer-lived winter annuals, germinating in the fall and flowering in early spring (Wilczek et al., 2009). Experiments have shown that winter cold is a major factor limiting performance (Ågren & Schemske, 2012; Korves et al., 2007). Additionally, Arabidopsis lacks physiological traits for dealing with severe water deficit so drought likely limits individual performance in nature (Clauw et al., 2015). Studies of large-scale environmental response in Arabidopsis have focused on the role of local adaptation in genetic diversity in the species (e.g. Hancock et al., 2011; Lasky et al., 2018; Martínez-Berdeja et al., 2020; Toledo et al., 2020), but less is known about the determinants of the species’ distribution. The last overview of the climate biogeography of Arabidopsis was Hoffmann (2002), who considered Arabidopsis native to western Eurasia, but recently introduced in China and much of Africa. However, studies show these latter populations are old, being genetically diverse, and with many unique genetic variants (Durvasula et al., 2017; Zou et al., 2017).

Distributions are dynamic through time due to environmental change, dispersal, and demographic stochasticity. Studies have used genetic data to infer Arabidopsis refugia during the last glacial maximum (LGM), where populations (sometimes referred to as “relicts”) persisted locally before subsequent admixture with an expanding, now widely distributed “non-relict” lineage (Lee et al., 2017). Whether these refugia corresponded to suitable climates is less clear. Existing projections of future climate impacts on Arabidopsis have focused on *relative* climate impacts on different genotypes (Exposito-Alonso et al., 2018; Fournier-Level et al., 2016), rather than distribution dynamics. Additionally, many species have colonized new regions due to human introduction, sometimes exhibiting traits different from native range populations, potentially in response to new environments (Turner et al., 2015). Arabidopsis has colonized many regions, but it is unclear to what degree these represent novel environments.

To evaluate biogeographic hypotheses linked to organismal and population processes, we used recently developed modeling approaches and updated climate and occurrence data. We combined occurrences (including many outside Europe poorly represented in previous work) with climate data to build distribution models, and then test how model predictions correspond to performance estimated from herbarium specimens and genetic variation in natural accessions. We ask the following questions:

1. What climate factors constrain the distribution of Arabidopsis? We hypothesize that winter cold and summer drought are the most important constraints, depending on region.
2. Are individual performance and genetic variation associated with model-estimated habitat suitability? We hypothesize that individuals reach larger sizes in regions with greater suitability and that populations adapt along gradients in suitability through changes in traits such as flowering time, a key component of life history.
3. Are occurrences outside the native range predicted by a native range model, suggesting stable realized niches following colonization? Or is there evidence Arabidopsis has colonized novel environments?
4. Where did Arabidopsis persist during the Last Glacial Maximum (LGM)? And where will Arabidopsis move in future climates?

## Methods

### Occurrence data

We developed a set of high-quality occurrence data (*i.e*. species ID verified and location checked, N=4,024) from published research (Durvasula et al., 2017; Hsu et al., 2019; Mandáková et al., 2017; Zacchello et al., 2022; Zeng et al., 2017; Zou et al., 2017), publicly available herbarium and germplasm accessions with known collection locations (Alonso-Blanco et al., 2016; DeLeo et al., 2020), and some of our recent field collections in East Africa (Gamba et al., 2022). These span a period of 1794 - 2018. The herbarium specimens and new collections include little-studied populations in Saudi Arabia, Somalia, Djibouti, Eritrea, Rwanda, Ethiopia, Uganda, Sudan, and Nepal. Duplicate occurrence points were eliminated (samples are often split and sent to different herbaria). For model fitting, we excluded occurrences from putative non-native regions (the Americas, New Zealand, Japan).

We also used occurrence data (with coordinates and without flagged problems N=115,226) from the Global Biodiversity Information Facility (GBIF) to test model predictions in regions outside of the native range of Arabidopsis (downloaded 30 Dec 2020, Gbif.Org, 2020). We deem these occurrences as lower quality given that many have not had the species identity and location checked (DeLeo et al., 2020).

### Environmental data

Climate data were extracted from CHELSA (Climatologies at High resolution for the Earth’s Land Surface Areas) v1.2 at 30 arc second spatial resolution (Karger et al., 2017). Current conditions are the average of 1979-2013 estimates. We selected the following climate variables based on hypothesized importance (Gienapp et al., 2017; Hancock et al., 2011; Lasky et al., 2014, 2018) and relatively low inter-correlation (Pearson correlation coefficients among variables at occurrences < 0.75): isothermality (Bio3), minimum temperature of coldest month (Bio6), temperature annual range (Bio7), mean temperature of wettest quarter (Bio8), mean temperature of the warmest quarter (Bio10), precipitation seasonality (Bio15), precipitation of wettest quarter (Bio16), and precipitation of driest quarter (Bio17). We also included elevation from Tozer et al., (2019).

For projecting past distributions, we obtained climate estimates from the last glacial maximum (LGM) at 21k yrs before present from CHELSA PMIP3 (Karger et al., 2017). We used the global elevation and bathymetry map with 15 arc second resolution from Tozer et al. (2019) and we adjusted elevations to make sea level 134 m lower than present (Lambeck et al., 2014) to project potential suitable habitat at the LGM on land in areas currently submerged. For projecting future distributions, we used climate projections from five divergent global climate models for 2050 from CHELSA v1.2 using the RCP 4.5 emissions scenario (Karger et al., 2017). We also show RCP 6.0 in the supplement for context (Figure S13), though it is highly similar to 4.5 in the target time period.

To characterize temporal variation in climate (climate anomalies), we used the Climate Research Unit (CRU) TS 4.01 dataset, providing a global time series of monthly temperature and precipitation for the period 1900-2010 at a 0.5° resolution (Harris et al., 2014). From the CRU data we calculated the same bioclimatic parameters that we used from CHELSA, but in the CRU data these bioclimatic variables were specific to each herbarium specimen in the time period it was collected (Supplemental Methods). We then calculated local anomalies for each of these variables by taking the observed value, subtracting mean across the entire time series, and dividing by the standard deviation (DeLeo et al., 2020).

### Performance estimates from herbarium specimens

We estimated fecundity on a subset of herbarium specimens using two traits. First, we measured the length of the longest inflorescence, reasoning that longer inflorescences would have more fruits and seeds. Second, we measured maximum rosette leaf length, reasoning that larger rosettes would support later reproductive investment if these collected plants were allowed to continue growth *in situ* (see Supplement for a validation). We used ImageJ on 2,194 specimen images to estimate the tallest point of each inflorescence (N=2,188) and the maximum rosette leaf length (N=1,264; see Supplement).

### Range-wide genetic variation in life history

Many late-flowering Arabidopsis genotypes require cold cues (vernalization) to flower and also show slower growth and more stress tolerance, delineating a life history axis (Lovell et al., 2013; Vasseur et al., 2018). To assess life history variation, we used published experimental data on flowering time for 898 whole-genome resequenced accessions from the native range with reliable geographic coordinates grown at 10 and 16°C (The 1001 Genomes Consortium 2016). To estimate vernalization sensitivity, we calculated the difference between flowering time at 10 and 16°C.

### Species Distribution Modeling

We thinned the original 4,024 high quality occurrence points to one sample per 1 degree grid cell to reduce sampling bias (N=764) using the ‘sp’ package (Bivand et al., 2008). To characterize potentially inhabited sites, we generated pseudoabsence background points using the ‘dismo’ package (Hijmans & Elith, 2013) by sampling 10,000 random points within a 500 km buffer around occurrence points. We limited this buffer to 500 km to avoid including low sampled regions in Africa and Asia that might result in biased model fit. We present a result with 1,000 km buffers in the supplement for reference.

We used Maxent version 3.4.0 to generate a species distribution model (Phillips et al., 2006). MaxEnt was implemented with the R package ‘ENMeval’ v2, and parameters were optimized using the ‘checkerboard2’ method for cross validation (Muscarella et al., 2014).

Among the tested settings (ENMeval defaults), we chose the model with lowest AICc value and used this to project habitat suitability under recent conditions across the globe. For all models we used the logistic output of MaxEnt that scales suitability from zero to unity.

We used permutation importances to determine which climatic factors drove predictions in the distribution model. We also used the ‘limiting’ function in the R package ‘rmaxent’ to determine the most limiting climatic factors in each location, defined as the environmental covariate that has the largest decrease in suitability at a given location relative to the suitability achieved if the covariate had its value equal to the global mean (Baumgartner et al., 2017; Elith et al., 2010). State another way, the local limiting factor is the environmental condition most limiting suitability, compared to an alternative scenario where that condition takes its global mean.

In non-native regions, we evaluated whether Arabidopsis is limited from further expansion at range edges by climate, *i.e.* whether there were no more unoccupied suitable environments near existing populations. To do so, we calculated suitability in a zone 50-100 km from existing GBIF occurrences and compared occupancy in these buffers in the native range to invaded regions.

We also projected the MaxEnt model using past (LGM) and future climate conditions.

For future conditions, we calculated the mean and standard deviation of habitat suitability projected for the five climate models. To assess whether model predictions were extrapolating into poorly characterized or novel climates, we compared the present-day model training climates to each predicted climate conditions, calculating multivariate environmental similarity surfaces (MESS) following Elith et al. (2010). Higher values on a MESS map indicate conditions in a location (or point in time) are similar to the reference environmental conditions used to fit the model. Negative values indicate that at least some variables are outside the range of environments used to fit the model, signifying extrapolation into novel environments (Elith et al., 2010).

### Performance and habitat suitability

We asked whether suitability corresponded to plant size. We first tested these relationships with Pearson’s and Spearman’s correlations. We also fit Generalized Additive Models (GAMs) where herbarium specimen sizes were the response variable. The model included covariates for suitability at the collection location (square-root transformed to reduce the lower-tail influence) and (as a nuisance variable) the year of collection (scaled to mean zero and unit variance) to account for potential changes in size over time. We used GAMs to allow smooth spatial variation in parameters, allowing us to capture regional variation in environmental response (for an extensive description of GAMs in biogeography see Guisan et al., 2002). The model can be represented as:

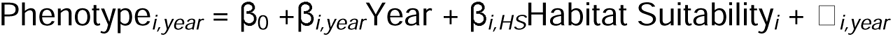

where *i* represents the specimen location. Errors were modeled as normally distributed. Spatially-varying parameters were constrained to smooth spatial variation in the GAMs using the ‘mgcv’ package for R, which we used with REML to fit all GAMs (Wood, 2011). We considered covariates to be significant at a given location if their 95% confidence interval (CI) excluded 0, and the Moran’s I (Cooper, 2021) for model residuals did not indicate spatial autocorrelation.

We also asked whether local limiting factors corresponded to climate variables with significant effects on plant performance. We hypothesized that for a given region, temporal fluctuations in local limiting factors would be associated with temporal variation in plant size in herbarium specimens. To address this hypothesis, we tested if specimen sizes were correlated with anomalies in climate conditions corresponding to the climate variables used in the MaxEnt model. We used the yearly climate anomalies we calculated from CRU in GAMs with spatially varying coefficients, where for a phenotype measured at location *i*:

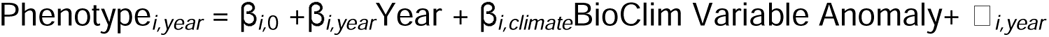

We then qualitatively assessed whether local limiting factors inferred by MaxEnt had anomalies that were correlated with plant size (local β*_i,climate_*coefficients in the GAMs).

### Genetic variation and habitat suitability

We tested how several aspects of genetic variation are related to suitability. First, we assessed whether there was a genetic signature to genotypes currently found in regions that were also highly suitable during the LGM. Lee et al. (2017) identified outlier haplotypes across the genome that were highly differentiated, hypothesizing that these were genomic regions deriving from long isolated “relict” populations that persisted through glacial cycles. We used bivariate correlations and Fisher’s exact test to test if higher LGM suitability was associated with a greater number of outlier genomic regions.

Next, we focused on genetic variation in a key life history trait, flowering time. We estimated linear models relating suitability and flowering time. To account for population structure and neutral processes that can affect spatial variation in flowering time, we also tested whether suitability was associated with flowering time after using random effects for genome-wide similarity among accessions. A significant suitability-flowering time association in this model would suggest selection linked to suitability acts on flowering time. This test is akin to *Q_ST_-F_ST_* contrasts (Whitlock & Guillaume, 2009), except that an explicit environmental gradient is tested (suitability). We implemented this mixed model the function ‘lmekin’ from the R package ‘coxme’ (Therneau & Therneau, 2015), along with ‘kinship2’ (Sinnwell et al., 2014). The kinship matrix was obtained from 2,027,463 published whole genome resequencing SNPs (Alonso-Blanco et al., 2016). Additionally, to test for geographic variation in suitability-flowering time relationships, we fitted GAMs of flowering time with spatially varying coefficients:

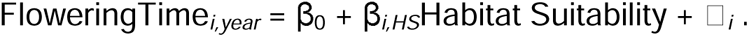

Our last strategy was to scan the Arabidopsis genome for genes where different alleles were found in high versus low suitability locations. This approach could be considered a novel application of a genotype-environment association (GEA) in an environmental genome wide association study (GWAS) (Lasky et al., 2023). Such a change in allele frequency across gradients in suitability would suggest that this genetic variation was involved in local adaptation to low versus high suitability environments. We used univariate linear mixed-effects models in GEMMA (v 0.98.3) (Zhou & Stephens, 2012) to perform the environmental GWAS in a set of 2,053,939 SNPs filtered for MAF=0.05 from 1003 native-range ecotypes part of the 1001 genomes panel. GEMMA includes random effects that are correlated according to genome-wide similarity between samples (“kinship”), allowing it to account for genomic background effects.

## Results

### Q1. Climate constraints on the distribution of Arabidopsis

Our optimization of MaxEnt models (with AICc) resulted in a selection of a model with linear and quadratic effects, with the regularization multiplier of 0.5. The training AUC value was 0.79 and the average test AUC with checkerboard2 cross validation was 0.78. The omission rate of the 10th percentile of suitability for training points was 0.10, suggesting our models were not overfit as they were able to predict low probability occurrences as well as expected (Fielding & Bell, 1997; Peterson et al., 2011). The areas of high suitability overall correspond well to the documented Arabidopsis distribution, with one notable exception being tropical lowland sites (Congo basin) in sub-Saharan Africa (Figure 1A). This region was near zero multivariate environmental similarity to training data, indicating the model may have been poorly constrained there, while most of the regions of high predicted suitability where Arabidopsis is documented have positive similarity (MESS, Figure S2).

**Figure 1.**
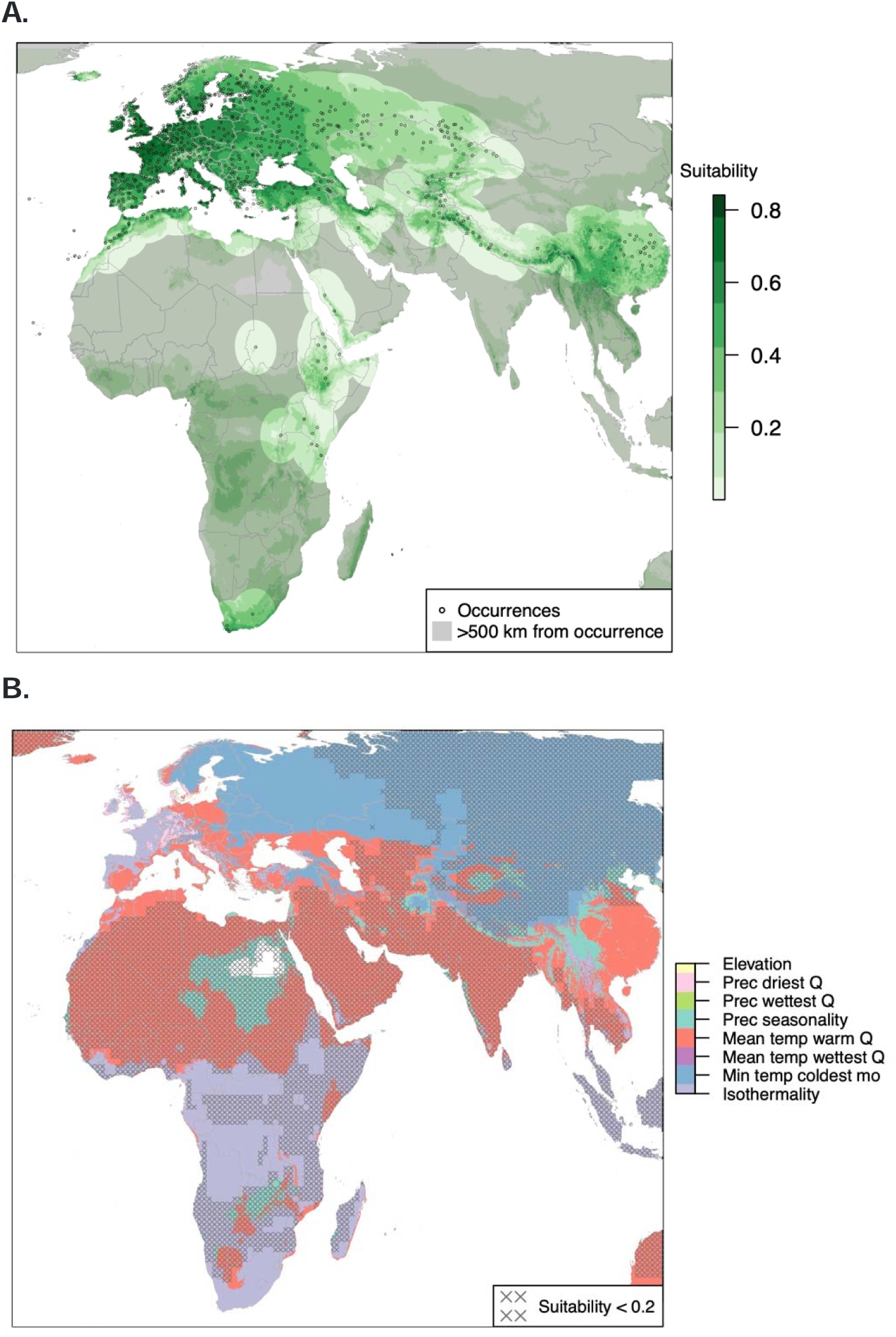
(A) Habitat suitability (green) under current climate conditions with thinned occurrences (n = 662) used in fitting shown as black circles and (B) limiting factors from the MaxEnt model fit to current climates. Regions far (>500 km) from known occurrence have a gray mask in (A). For reference, regions of very low suitability (less than 0.2) in (B) are marked with gray ‘x’ symbols. Equal Earth projection was used. Abbreviations in (B) as follows: precipitation of the driest quarter “Prec driest Q”, precipitation of the wettest quarter “Prec wettest Q”, precipitation seasonality “Prec seasonality”, mean temperature of the warmest quarter “Mean temp wrm Q”, mean temperature of the wettest quarter “Mean temp wettest Q”, temperature annual range “Temp ann range”, minimum temperature of the coldest month “min temp coldest month.

The permutation importance of the model covariates across the entire native range revealed that the minimum temperature of the coldest month (PI = 38%) and the mean temperature of the warmest quarter (PI = 31%) were the two most important variables, suggesting winter cold stress and summer heat stress are most important in constraining Arabidopsis’s distribution (Table S1). The next most important variable was isothermality (PI =15%), as Arabidopsis tends to be found in regions with low isothermality (e.g. most temperate zones).

To identify spatial variation in climate constraints, we also identified local limiting factors (Figure 1B). Across northern Eurasia, minimum temperature of the coldest month limited habitat suitability. Across the Mediterranean and tropical/subtropical regions, the temperature of the warmest quarter was limiting. Across Eastern Europe and Central Asia, winter cold was limiting adjacent to other regions where summer heat was limiting, highlighting the multiple stressors in this region.

We hypothesized that for a given region, temporal fluctuations in local limiting factors would be associated with temporal variation in plant size in herbarium specimens. We found that years with warmer minimum winter temperatures were associated with significantly taller inflorescences from Turkey to central Asia (Figure S3, panel B). Concordantly, in much of this region the Maxent model indicated that minimum temperature of the coldest of the month was the most limiting factor (namely in the Caucusus and from Kazakhstan to northern India, Figure 1B). We also found that years with greater seasonality of precipitation were associated with shorter inflorescences in central Asia, but taller inflorescences in central Europe (Figure S3, panel F). Partly consistent, in the southern part of this central Asian region the MaxEnt model indicated precipitation seasonality was limiting (Figure 1B). Years with higher isothermality were positively associated with inflorescence height in Eastern Europe but this was not a limiting factor in this region (Figure S3, panel A). Other climate anomalies were not significantly associated with temporal variation in plant size (Figures S3 and S4).

### Q2. Declining individual performance and prevalence of stress escape life history in low suitability regions

As habitat suitability increased, so did the inflorescence height of individual plants in herbarium specimens (Pearson’s r = 0.12, p < 10^-8^; Spearman’s rho = 0.10, p < 10^-6^; n = 2169). The relationship with suitability was even stronger for maximum rosette leaf length (r = 0.25, p < 10^-16^; rho = 0.26, p < 10^-16^; n = 1253, Figure 2), which was also more correlated with total fruit + flower number (Supplement), suggesting fecundity was greater in regions of high predicted suitability. We tested size-suitability relationships using GAMs with spatially varying suitability coefficients (but not including climate anomalies – distinct from the previous section Q1). We found a consistently positive relationship between suitability and size that was significant for rosette leaf length in western and central Europe and for inflorescence height across most of Eurasia (Figs S5 and S6). Unexpectedly, in this model where year of collection was considered primarily as a nuisance variable, inflorescence height significantly declined over time in northwest Europe (Fig S6; e.g. in Scandinavia and Scotland, maximum rosette leaf length vs year, Spearman’s rho = -0.14).

**Figure 2.**
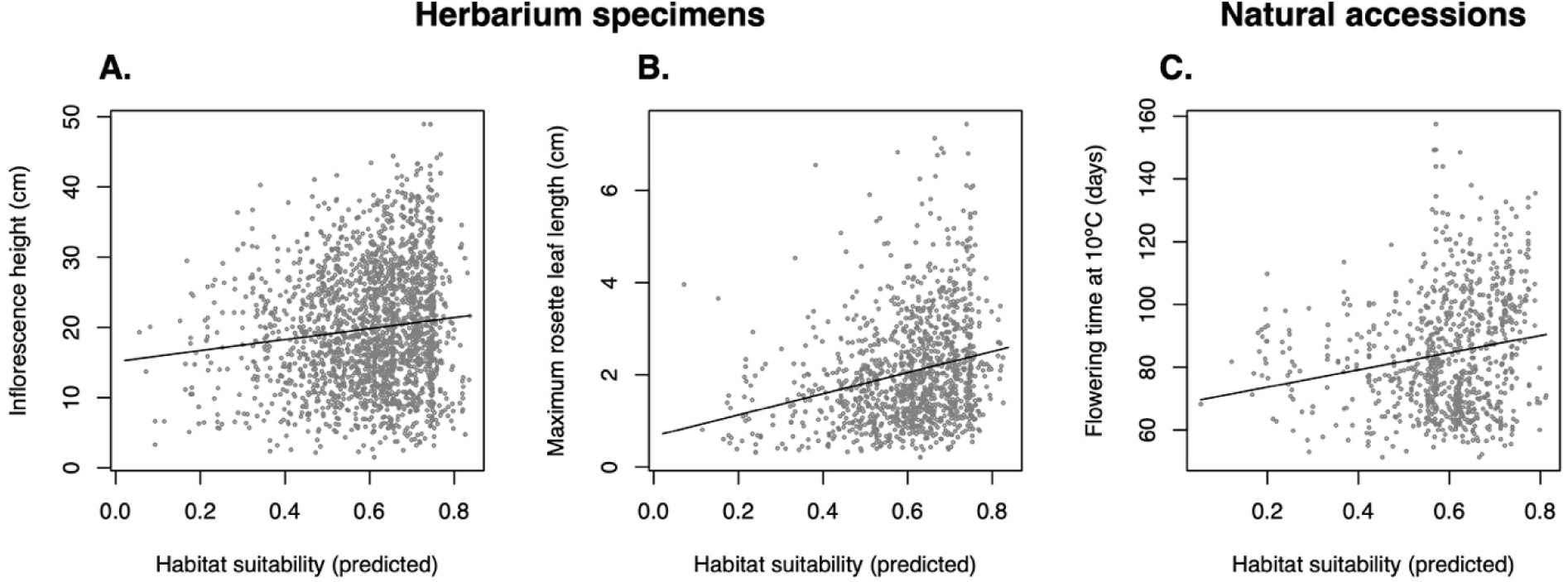
Predicted current habitat suitability compared with individual plant size (A & B) and genetic variation in a measure of life history (C). Size measures include inflorescence height (A, relationship with suitability: Pearson’s r = 0.12, Spearman’s rho = 0.10) and maximum rosette leaf length (B, r = 0.25, rho = 0.26) from herbarium specimens. Flowering time of natural accessions (C, r = 0.21) was taken from published data on a growth chamber experiment at 10°C (Alonso-Blanco et al., 2016). Linear model fits are shown. N = 2053 for inflorescence height, N = 1179 for maximum rosette leaf length, and N=953 for flowering time.

We compared habitat suitability with published data on genetic variation in flowering time. We found that flowering time at 10°C and flowering time plasticity (flowering time at 10°C – time at 16°C) were significantly positively associated with suitability (r=0.19, p<10^-8^ and r=0.13, p=0.0002, respectively), but not for days to flower at 16 °C (r=0.05, p=0.1264, Figures 2 & S7). In GAMs with spatially-varying suitability coefficients, the suitability-flowering time pattern were largely consistent across Eurasia (Figure S8). The suitability-flowering time association was significant even when accounting for genomic similarity among accessions, suggesting selection associated with suitability could maintain geographic clines in flowering time. Specifically, linear mixed-effects models found a positive association for flowering time at 10°C (p=0.0004, n=884), and also at 16°C, the latter of which may have been obscured by population structure that was unaccounted for in the simple linear regression model (p=0.02, n=852, Tables S2-S3). The suitability association was not significant for plasticity in flowering time when accounting for genomic similarity (p=0.77, n=852) (Table S4). Consistent with this potential obscuring of population genetic structure, we found that 10 ADMIXTURE genetic clusters in Arabidopsis (Alonso-Blanco et al., 2016) were significantly different in their habitat suitability (F(9,993)=201.6, p<10^-16^, Table S5, Figure S9).

We scanned the genome for genes that showed allele frequency correlations with suitability. The most strongly associated SNP was in the putative promoter region (134 bp from the start, linear mixed model, p < 10^-16^) of ERF53 (AT2G20880), a transcription factor that regulates response to drought, salt, and heat (Figure S11) (Cheng et al., 2012; B. Li et al., 2019). This SNP showed a strong allele frequency cline from Europe to Asia, where the alternate allele was nearly fixed in accessions east of the Ural Mountains, which is a region of low estimated suitability (Fig S10). Furthermore, the alternate allele was associated with lower expression of ERF53 (Wilcox test, p = 0.0013) in published transcriptome data (Kawakatsu et al., 2016), suggesting a *cis-*regulatory variant is locally adapted to low suitability parts of Asia. Relatedly, Jiang et al., (2022) showed that knockouts of ERF53 have temperature-dependent effects on fecundity.

### Q3. Using the native range model to predict outside the native range

Australia and New Zealand occurrences were largely in areas predicted to be highly suitable (Figure 3). By contrast, North America includes occurrences in highly suitable areas (Pacific Northwest) but many in low suitability areas (interior east, where summer heat was predicted limiting, Figure S12). The eastern US also has near zero similarity on the MESS map, suggesting a novel set of conditions compared to the native range (Figure S2). Similarly, the regions where Arabidopsis occurred in Korea and Japan were of lower suitability than in the native range, also with summer heat as the predicted limiting factor (Figure S12), suggesting Arabidopsis in eastern North America and East Asia inhabits climates with distinctly warm summers.

**Figure 3.**
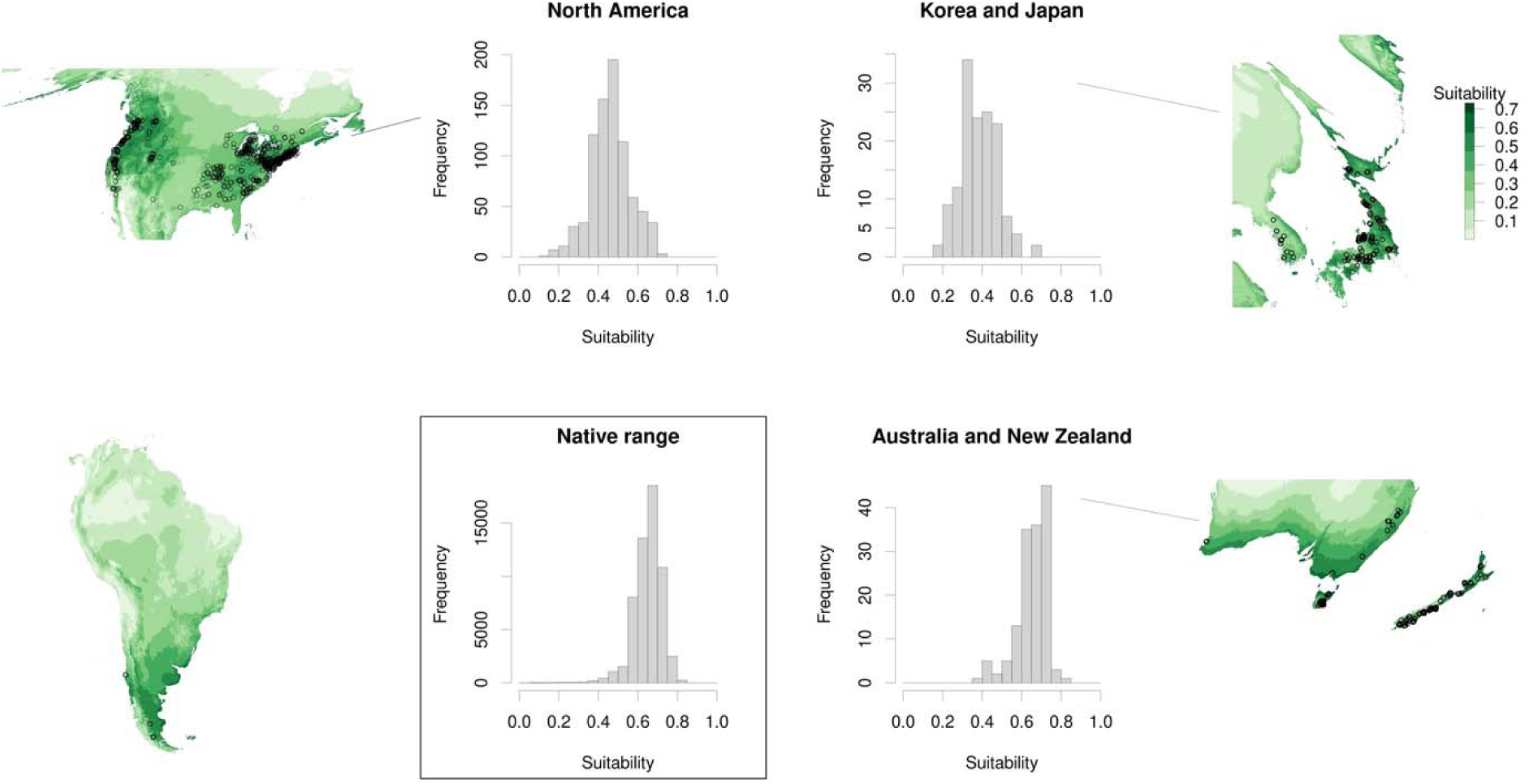
Distribution of predicted habitat suitability (based on our native-range MaxEnt model, underlying map surfaces in green) for GBIF occurrences in various regions (black circles). We do not include a histogram for South America because there are too few occurrences. Equal Earth projection was used.

We investigated whether non-native Arabidopsis occupy all available habitat in their regions or whether suitable habitat remains unoccupied. We found that locales 50-100 km from occurrences included some areas of high suitability in North America, suggesting suitable sites remain unoccupied (4.3% of grid cells in these 50-100 km regions had suitability > 0.6). There were extensive areas of high suitability along the Pacific coast of North America to around 60°N, but no occurrences north of 50°, which we confirmed with an expert botanist (pers comm.

Matthew Carlson). The timing of initial colonization is likely not a factor given that occurrences from near Vancouver date at least to 1939. This region has near zero similarity on the MESS map, potentially indicating that the native range model is not well constrained there (Figure S2). Similarly, the southern coast of Australia is highly suitable but Arabidopsis is apparently absent (35.0% of these 50-100 km regions with suitability > 0.6). Records are restricted to the southeast and southwest (confirmed by botanists, Shelley James, Tim Entwistle, Neville Walsh pers. comm.), even though records date at least to 1959 in SE Australia. In South America there are a few Arabidopsis records from the Southern Cone, and it appears to be rare in the region (pers. comm. Diego Salariato), while apparently suitable environments occur throughout Patagonia and high elevation Andean sites that are apparently unoccupied (pers comm. Gwendolyn Peyre, Santiago Madriñán). Large areas of southern Australia and South America show positive similarity on the MESS map suggesting the model is well constrained in those regions. By contrast in Korea and Japan there are very few sites expected to be highly suitable that are not already occupied by Arabidopsis (2.4% of these 50-100 km regions with suitability > 0.6).

### Q4. The distribution of Arabidopsis during the Last Glacial Maximum and in the future

We projected suitability onto the climate at the LGM and found several putative refugia, where past environmental conditions could have supported Arabidopsis persistence. In particular, the Mediterranean/Caucuses/south Caspian Sea, larger portions of Africa > 1000 m (current) asl, and China and SE Asia appear as refugia (Figure 4). North Africa, the Atlantic European coast, and the islands of Sicily, Corsica, and Sardinia also appear as highly suitable potential refugia.

**Figure 4.**
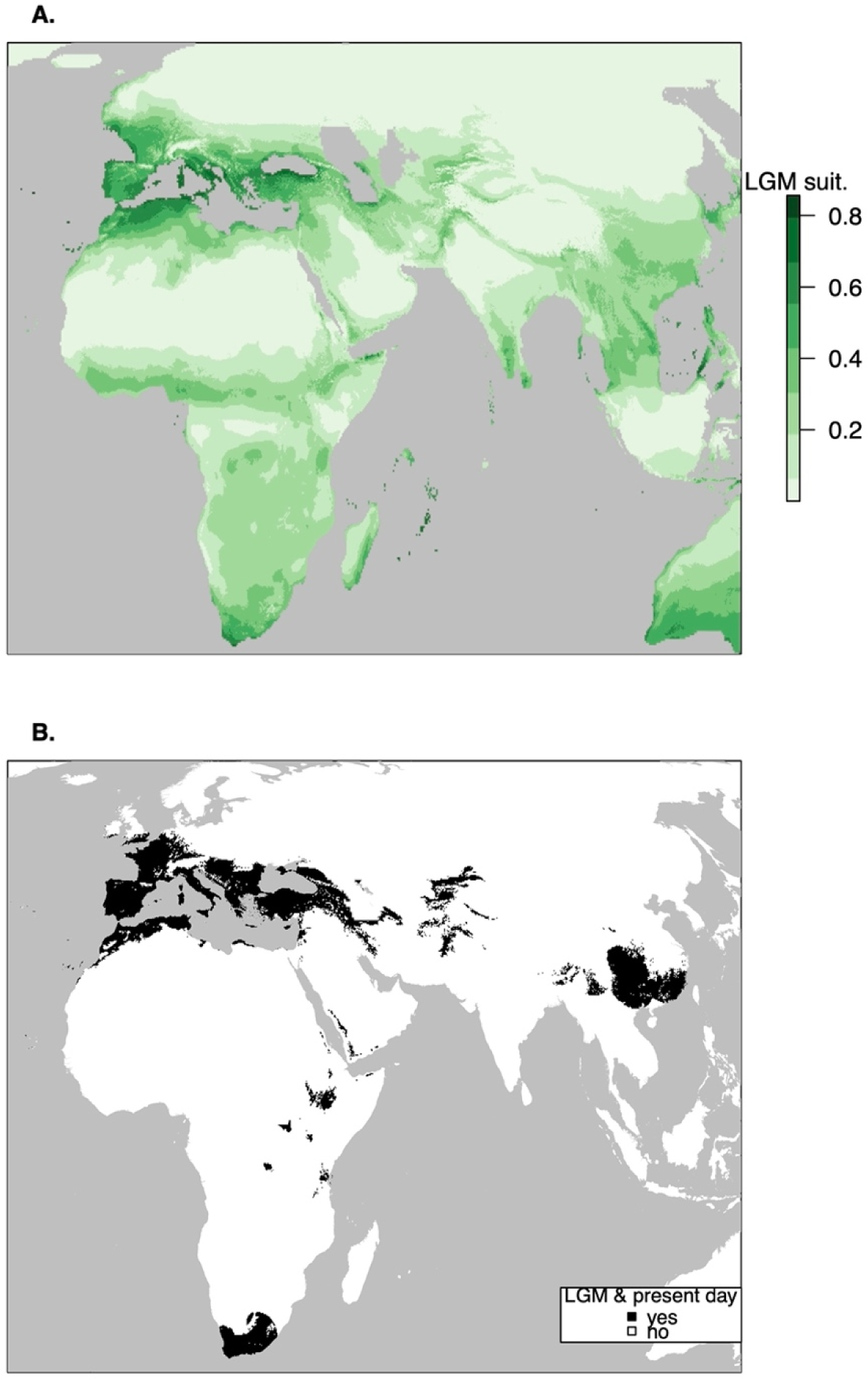
(A) Predicted distribution during the Last Glacial Maximum and (B) areas in black with suitability > 0.25 during both the LGM and current conditions as well as within 500 km of known current occurrences. Because of their greater level at the LGM, in (A) we masked the LGM Caspian and Aral Seas (including regions of high putative suitability) from the map (Prentice et al., 1993). Lake Victoria was left unmasked as it was likely very low during the LGM (Johnson et al., 1996). Equal Earth projection was used. For (B), the threshold of 0.25 was chosen for reference as it approximately matches the current distribution of Arabidopsis; the alternate thresholds for visualization of 0.2 and 0.3 and be found in Figures S14 and S15.

We projected climate suitability for Arabidopsis in the year 2050. Some current high suitability regions will remain so, such as in Northern Europe. Nevertheless, we found poleward and up-elevation shifts in regions of high suitability and retreats at lower latitudes and elevations. In the native range, all African sites show declining suitability, as did most Mediterranean sites, highlighting vulnerability of these populations. By contrast, Arctic Europe, and the mountains of central Asia/Tibetan plateau show increased suitability, highlighting potential range expansions (Figure 5).

**Figure 5.**
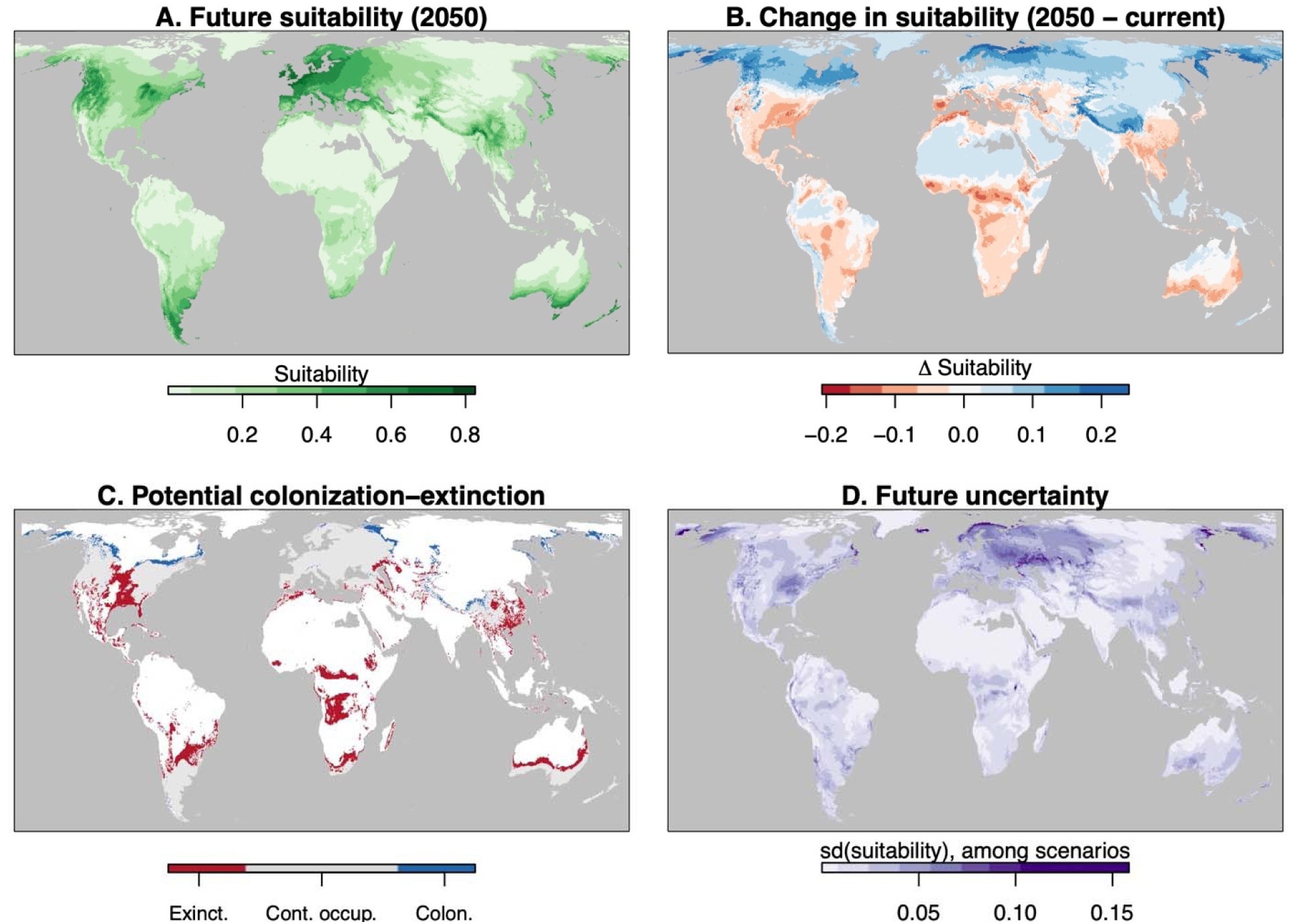
Future predicted suitability for Arabidopsis (A), change in suitability from future compared with present (blue indicates improving suitability and red decreasing, B), regions of potential colonization (blue) continued occupancy (gray), and extinction (red) based on a threshold suitability of 0.25 for occupancy (C), and the standard deviation in suitability among the 5 tested climate models giving uncertainty (D). Equal Earth projection is used. RCP 4.5 emissions scenario is shown, see Fig S13 for highly similar patterns under RCP 6.0. For (C), the threshold of 0.25 was chosen for reference as it approximately matches the current distribution of Arabidopsis; the alternate thresholds for visualization of 0.2 and 0.3 and be found in Figure S16 and S17.

We found concordance between our LGM predictions and the frequency of outlier genomic regions in modern genotypes (Lee et al., 2017). The suitability during the LGM at the site of genotype origin was positively correlated with the number of outlier haplotypes they carried (N = 933, Pearson’s r = 0.22, p<10^-12^; Spearman’s rho = 0.14, p<10^-5^). This relationship was partly due to the fact that >90% of the distinct relict genotypes defined by Lee et al. (2017) were located in regions where the LGM suitability was over 0.25, versus only 48% for non-relicts (Fisher’s exact test p<10^-5^, 0.25 approximately defining range limits in current climates, but this test was highly significant for suitability thresholds from 0.1 to 0.5). However, even among non-relicts, the higher the LGM suitability the greater the frequency of outlier haplotypes (N = 911, Pearson’s r = 0.20, p<10^-10^; Spearman’s rho = 0.10, p=0.0017).

## Discussion

We used the model plant Arabidopsis in a case study of integrative climate biogeography of a species’ past, present, and future distributions. This species is key to a large body of plant biology research, and an in-depth study connecting its biogeography to genetic and phenotypic variation provides important context for the biology of Arabidopsis. The size of individual plants in herbarium specimens, as well as genetic variation in flowering time and a gene controlling abiotic stress response, suggested that regions with lower predicted suitability harbored populations with altered life history and physiology. The consistency between model predictions and individual variation provides partial model validation, bolstering our confidence in conclusions from model projections. Using these model projections, we identified new glacial refugia, and looking to the future, we found that genetically distinct lower latitude populations are most threatened by climate change.

### The current distribution of Arabidopsis and its limiting factors

The fitted native-range model largely corresponded to occurrences and indicates Arabidopsis broadly distributed across Europe, moister regions of central and eastern Asia, and mountains across Africa. Our results advance beyond the most recent climate biogeographical study of Arabidopsis by Hoffmann (2002), who used monthly climate data (but notably no synthetic bioclimatic variables) from Leemans & Cramer (1991) mostly to qualitatively describe the regions occupied by Arabidopsis. Our study included populations in sub-Saharan Africa and Asia that were overlooked or considered non-native by Hoffmann (2002). We also included occurrences from regions apparently missed by Hoffmann (2002). Zou et al. (2017) more recently built models of Arabidopsis’s distribution using Eurasian populations and default MaxEnt settings, but did not include occurrences from Arabia, sub-Saharan Africa, and much of the Himalayas, Russia, and central Asia, and did not subsample occurrence to reduce bias.

Likely as a result of these issues, the predictions of Zou et al. (2017) show a pronounced peak in suitability in Germany, which was densely sampled in the genomics studies used for occurrences by Zou et al. (2017), but predicted low suitability in much of the core European range and near zero suitability in sub-Saharan Africa, Arabia, and most of the Russian part of the species range.

The two dominant limiting factors in our model were winter cold (at higher latitudes and elevations) and summer heat (at lower latitudes and elevations). Winter cold is recognized to limit Arabidopsis performance, in particular winter cold appears to be a dominant force in local adaptation of Arabidopsis (Ågren & Schemske, 2012; Gienapp et al., 2017; Monroe et al., 2016). In southern Europe, where summer heat was inferred to be limiting, Arabidopsis flowers in early spring (DeLeo et al., 2020) and thus summer heat is not usually directly experienced. In spring in these regions, warm temperatures might not reach consistently stressful levels (e.g. to induce fruit abortion) (Warner & Erwin, 2005) but it may be that moisture deficit driven by evaporative demand is directly limiting in late spring. Where suitability was highest for Arabidopsis, including Britain and the Atlantic coast of France, high isothermality and high precipitation of the driest quarter were identified as limiting. However, interpreting limiting factors in an area of extremely high suitability (near one) is not meaningful given that suitability can scarcely be increased.

Despite inference of winter cold and summer heat as primarily limiting, these were only partly reflected by temporal fluctuations in individual plant performance from herbarium specimens. Winter cold and precipitation variability anomalies were significantly associated with specimen size in much of Asia, where these were also the MaxEnt modeled limiting factors from Iran and Kazakhstan to Afghanistan and the Himalayas, suggesting these climate factors truly limit Arabidopsis populations. However, there were discrepancies between individual performance and limiting factors, likely for several reasons. First, the size of specimens is an imperfect fitness proxy. Usually only reproductive individuals are collected in herbaria, and individuals that would die before reproduction are excluded. Second, individuals are not randomly sampled from populations (Daru et al., 2018). Third, MaxEnt models face limitations due to misspecification, problems with occurrence data, or a mismatch between covariates and the true ecological factors limiting populations. Nonetheless, the limiting factors we identified here fit well with our knowledge of Arabidopsis ecophysiology and natural history.

### Habitat suitability, individual performance, and life history

We found a decrease in plant size with decreasing habitat suitability across the range of Arabidopsis, suggesting that our model suitability captured a substantial part of environmental effects on individual fitness. Similar to our findings, Karasov et al., (2022) found declines in the size of natural Arabidopsis plants at latitudinal extremes of the European range. Given the great variability in individual plant performance within populations obvious to casual observers, it is unsurprising that suitability here explained a minority of total variation in individual size, leaving most variation unexplained (Figure 2). Furthermore, fecundity and fitness response to treatments often have low heritability in many species (Price & Schluter, 1991), even in controlled experiments (Lasky et al., 2015).

There have still been few studies of individual performance with the geographic scope that allows inference across a species range (Angert & Schemske, 2005; Csergő et al., 2017; Greiser et al., 2020; Samis & Eckert, 2007). In a synthesis of studies of 40 species, Lee-Yaw et al. (2016) found that, on average, individual performance and distribution-model inferred suitability decline beyond range margins. From 42 studies Lee-Yaw et al. (2021) found that 38% identified some predictive ability of distribution models for individual performance. However, many previous studies relied on intensive observations of a small number of populations, while our estimates of performance from thousands of herbarium specimens allowed us to cover most of the species range. The increased availability of digitized museum specimens with trait data indicates an opportunity to estimate performance across distributions for many species (Bontrager & Angert, 2015).

We found that low suitability regions had earlier flowering genotypes. This relationship was noisy, with early flowering genotypes frequent in all levels of suitability but later flowering genotypes lacking from the least suitable regions. The suitability-flowering time associations were significant when accounting for genome-wide similarity between accessions, suggesting they reflect selection associated with suitability. We interpret the direction of the relationship as indicating that when suitability is low, Arabidopsis employs stress escape strategies, *i.e*. a rapid life cycle during favorable conditions (Ludlow, 1989). Nevertheless, our findings suggest that low suitability central Asian populations have some stress tolerating mechanisms, as they harbor distinct allele at a transcription factor (ERF53, AT2G20880) that regulates response to abiotic stressors (Cheng et al., 2012; B. Li et al., 2019) including performance response to temperature (Jiang et al., 2022). The restriction of late flowering genotypes to more suitable regions is counterintuitive given physiological work showing these are more stress tolerant (Lovell et al., 2013). However, it may be true that more favorable conditions make possible a slow growing, slow flowering, freezing-tolerant winter annual strategy.

### The distribution of Arabidopsis outside its native range

Arabidopsis has spread across the globe, largely to climates well-predicted by native range models (western North America, Australia, and New Zealand), suggesting some stability in climate niche, but also to some regions predicted to be less suitable (eastern North America, Korea, and Japan). Whether these are truly of lower suitability is unknown without performance data. It may be that Arabidopsis has colonized novel environments in these regions, or that the low suitability is only a model artefact. Studies often find limited transferability among native and introduced range models (Early & Sax, 2014; Liu et al., 2020).

### The distribution of Arabidopsis at the last glacial maximum

Lee et al. (2017) hypothesized five glacial refugia for Eurasia to be in Iberia, Sicily, Balkans, the Levant, and Turkmenistan. These are consistent with our range reconstruction, although we do not find clear barriers of unsuitable climates that would have isolated Italian, Balkan, and Levant LGM populations. Furthermore, we note that Sardinia/Corsica appears as a refugium, and we note that narrow, highly suitable areas along the southern and eastern Caspian Sea could be the location of the hypothesized Turkmenistan refugium. Lee et al. (2017) further hypothesized that the bulk of current European genotypes derive from an expansion originating on the northwest coast of the Black Sea, where we found a strip of highly suitable conditions for Arabidopsis during the LGM. Additional genetic studies hypothesized a long history of existence in sub-Saharan Africa (Durvasula et al., 2017) and conditions during the LGM suggest the species could have been more widespread than currently. Concordantly, Chala et al., (2017) used models of the distribution of Afroalpine habitat generally during the LGM to find expanded areas in East Africa compared to present day, though some locations such as Jebel Marra, Sudan were still surrounded by landscapes of unsuitable habitat even at the LGM.

Our findings also may help explain the history of Cape Verde Island populations, which Fulgione et al., (2022) estimated were bottlenecked and split from a source population 5-10 kya. The source population appears to be extinct and but is most similar to current Moroccan populations (Fulgione et al., 2022). Currently there are no known African Atlantic coastal, low elevation populations that would have been sources. However, we found that during the LGM and presumably for some time after, low elevation coastal high suitability habitat extended as far south as Ras Nouadhibo on the African coast, suggesting a source for the Cape Verde immigrants.

We found that the North Atlantic European coast was highly suitable (including areas currently inhabited) during the LGM, but current British Isle and French populations do not show genetic signatures of refugia (Lee et al., 2017). In both cases, local, distinct genotypes that survived the LGM may have gone extinct following expansion by the now dominant European genetic cluster (“non-relicts”) (Fulgione & Hancock, 2018; Lee et al., 2017). The previous Arabidopsis distribution modeling of the LGM (Zou et al., 2017) did not identify France and NE Iberia as highly suitable during the LGM and did not mask the Caspian Sea (thus overinflating that refugium). Zou et al. (2017) also did not identify the African refugia (except for a small region in SE Africa) and potential refugia in mountains of SE Asia.

### The future distribution of Arabidopsis

Over the next decades conditions are predicted to worsen for Arabidopsis across large areas of its range. Central Spain and mountains in Africa and Arabia may see the worst changes. Impacts may already be emerging: our intensive field search in the Dai Forest of Djibouti in 2018 failed to yield any Arabidopsis, despite the presence of abundant (but likely more drought tolerant) annual mustards (Brassicaceae) *Erucastrum arabicum* and *Sisymbrium erysimoides* in otherwise appropriate habitat for Arabidopsis. Arabidopsis was collected in this juniper woodland in 1954 but recent decades have seen increased drought and tree mortality (Witsen, 2012), suggesting potential extinction of this isolated marginal Arabidopsis population. Given the unique genetic diversity of the model plant Arabidopsis housed in its lower latitude populations (Durvasula et al., 2017; Gamba et al., 2022; Hsu et al., 2019; Lee et al., 2017; Zou et al., 2017) the conservation of these populations could benefit plant biology research.

By contrast, conditions are expected to improve for the northernmost populations in Europe, suggesting a potential current colonization front of Arabidopsis, in addition to higher elevation locations in Tibet, the Caucuses, Ural, Alps, and Hengduan mountains adjacent to currently inhabited regions. Similar currently unoccupied higher elevations are available in many African mountains though these are usually small mountaintop areas and may be poorly characterized by CHELSA climate data (Karger et al., 2017). Future distribution models would benefit from improved environmental data for these high elevation tropical sites.

## Conclusion

Biogeographic patterns emerge from processes affecting individuals and populations. Species distribution models provide potentially powerful windows into past, present, and future macroecology, but they are rarely confronted with range-wide individual performance data. Here we showed that lower suitability habitats inferred from distribution models had smaller plants with distinct life history, suggesting a stress escape strategy. While the relationships were noisy it may still be remarkable that they emerge against the many microsite contributors to individual level variation in a habitat generalist annual plant. Arabidopsis populations are distributed across diverse climates, but genetically distinct populations in lower latitudes that are potentially valuable for research are also highly threatened by anthropogenic climate change in the next few decades. We believe that combining distribution models with individual data on genotype and phenotype across a species range can be a useful approach to dissect the organismal mechanisms underlying distributions and to validate distribution models.

## Acknowledgments

This manuscript benefited from the comments of three anonymous reviewers. Victoria Meagher and Jaden Hill measured morphology on herbarium specimens. Patrick Herné assisted in digitizing many herbarium specimens and Eugene Shakirov translated many Russian specimen labels. Data from the MNHN in Paris were obtained from the participatory science program ‘Les Herbonautes’ (MNHN/Tela Botanica) which is part of Infrastructure Nationale e-RECOLNAT: ANR-11-INBS-0004. This work was supported by NIH award R35 GM138300 to JRL. RM was supported by grant 2019-03758 from Vetenskapsrådet. We also thank volunteers for digitization and georeferencing through DigiVol, a website developed by the Australian Museum in collaboration with the Atlas of Living Australia.

## Conflict of interest

The authors declare no conflict of interest.

## Data availability

MaxEnt model projections, occurrence data, and herbarium specimen data will be stored in a Dryad repository upon acceptance of this manuscript. 1001 Genomes data were obtained from 1001genomes.org. Current and paleoclimate data were obtained from chelsa-climate.org. Time series climate data were obtained from data.ceda.ac.uk.

## Notes

### Competing Interest Statement

The authors have declared no competing interest.

### Summary of Updates

We have updated our framing of the manuscript. We have redone all analyses with a new version of the dataset.

