## Supplemental Material for "Climate biogeography of *Arabidopsis thaliana:* linking distribution models and individual variation"

**Methods**

We separately used five divergent global climate models from the fifth phase of the Coupled Model Integration Project (CMIP5) under the Relative Concentration Pathway (RCP) 4.5 emissions scenario: ACCESS-1-0, MIROC5, CESM1-BCG, CMCC-CM, and MPI-ESM-MR, and five models from the RCP 6.0 emissions scenario: CCSM4, CESM1-CAM5, CSIRO-Mk3-6-0 , FIO-ESM, and GFDL-CM3. These emissions scenarios are similar over our projected timeframe (2040-2060) and so we expect minor differences in predictions.

When calculating annual anomalies for bioclimatic variables, we used months we deemed most relevant to the herbarium specimen based on its date of collection. Specifically, for the minimum temperature of the coldest month, precipitation of the wettest and driest quarters, and precipitation variability, we used the 12 months preceding collection because we reasoned it would best capture these potential stressors. For the mean temperature of the warmest and wettest quarter we used the same calendar year of the collection, because we reasoned this would allow us to capture growing season conditions for the most relevant period for an individual. For temperature annual range and isothermality we used the fall (October) preceding collecting to the summer in the year of collection (September) to attempt to capture both the growing season of collection and its prior cool season.

To test if inflorescence height and maximum rosette length were good proxies for total number of fruits, we counted flowers and fruits using ImageJ on 311 broadly distributed specimens (Figure S1) from the *Muséum national d’histoire naturelle* (herbarium code P), which provided high-resolution scans, allowing us to resolve individual flowers and fruits. We found that inflorescence height and maximum rosette leaf length were positively correlated with total number of flowers and fruits (inflorescence height r = 0.46, n = 304; maximum rosette leaf length r = 0.50, n = 183).

When there were multiple plants on an herbarium sheet, we selected one based on tissue sampling for a previous study (DeLeo et al., 2020) or we selected one at random using a random number generator when there were fewer than 6 plants present. We did not measure inflorescence length on specimens where the main inflorescence was obviously bent or broken. Likewise we only measured maximum rosette leaf length for specimens where rosettes were preserved flat (as opposed to crumpled, folded, or broken).

For all GAMs, relationships were predicted over a 140 by 200 grid from -28.5 to 140.5° longitude and 23.9 to 71.0° latitude.

**Figure S1.** 311 specimens from the P herbarium (MNHN, Paris) for which we counted individual fruits and flowers to compare with inflorescence height and maximum rosette leaf length.

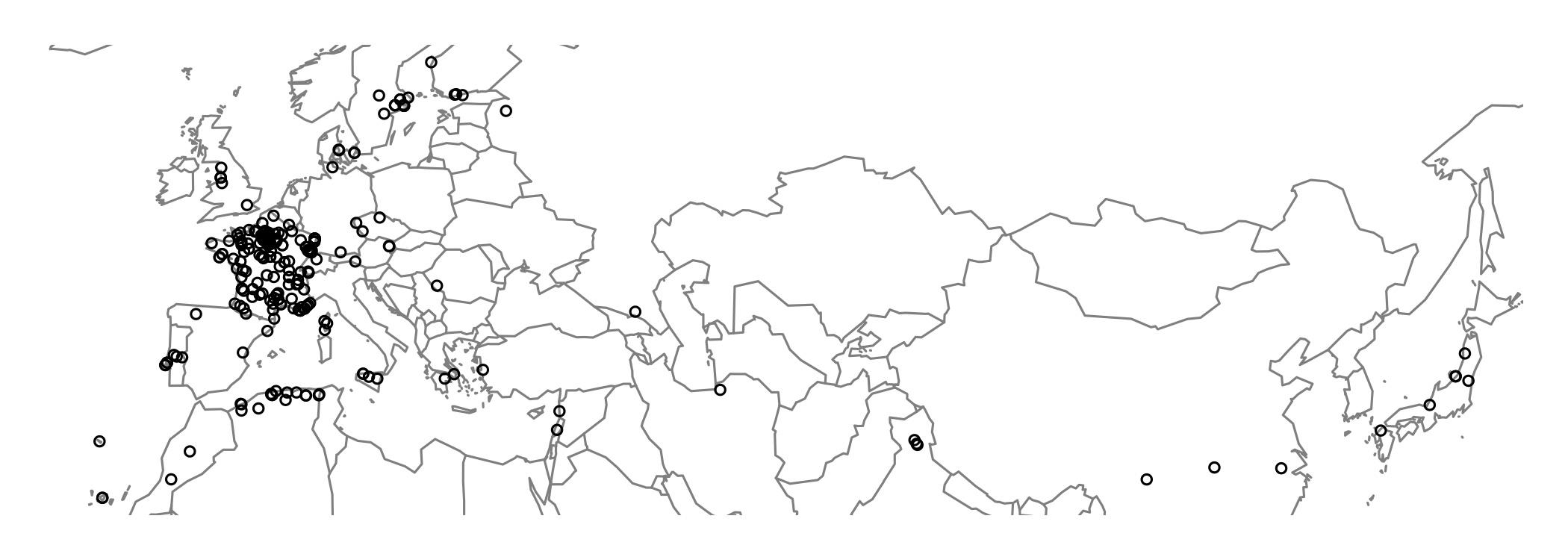

**Figure S2.** MESS maps of current, LGM, and future (RCP 4.5 mean) climate conditions. Blue indicates positive similarity and red indicates negative similarity with conditions used for fitting the model on present occurrences.

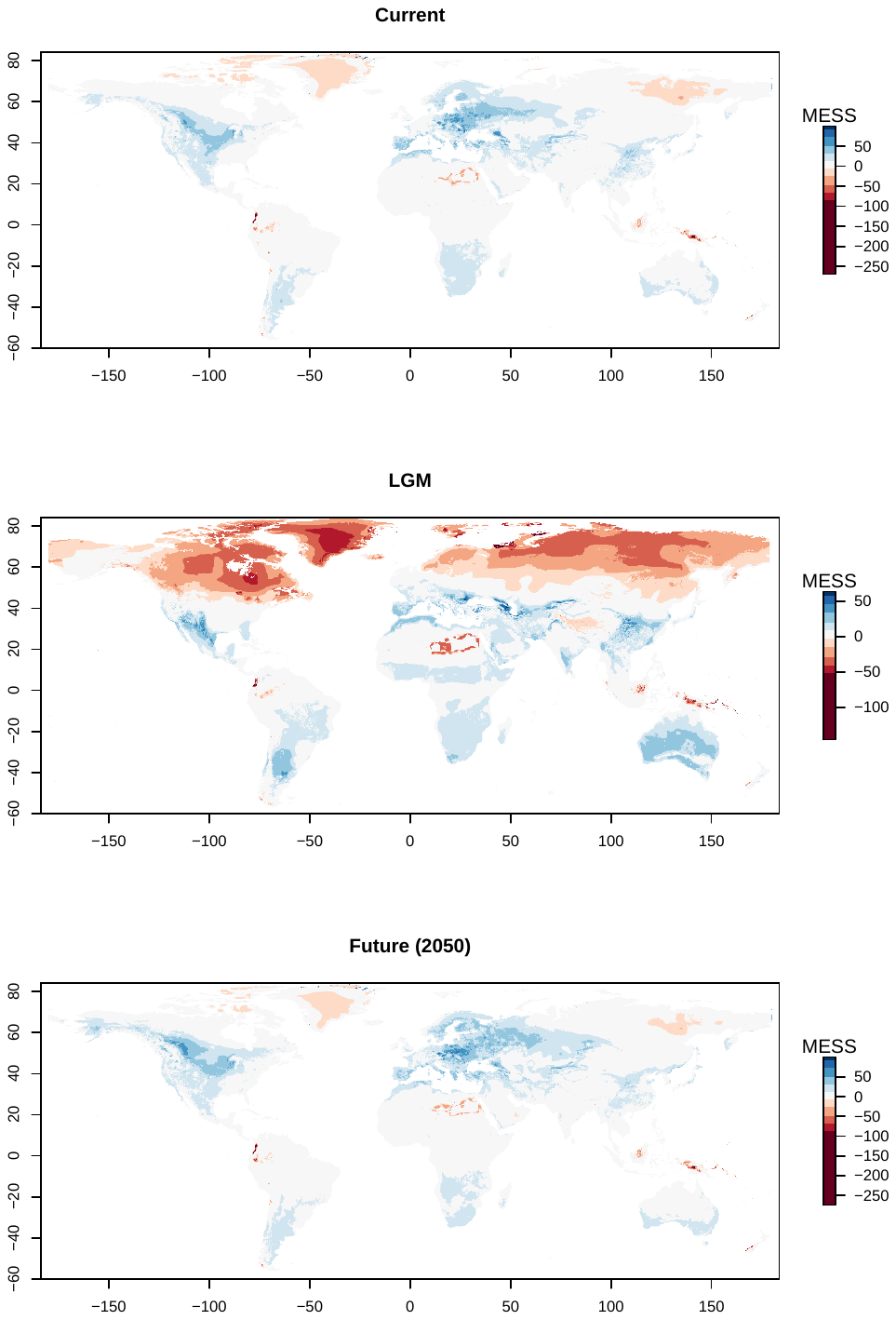

**Figure S3**. Spatial varying coefficients and intercepts from GAMs showing the relationship between inflorescence height and yearly anomalies in climate variables of isothermality (A, BIO3), mean daily minimum air temperature of the coldest month (B, BIO6), annual range of air temperature (C, BIO7), and mean daily mean air temperature of the wettest quarter (D, BIO8). Gray shading indicates areas where parameter values are not statistically significantly different from zero (e.g. western Europe in for the BIO3 effect in panel A), specifically where 95% confidence intervals overlap with 0.

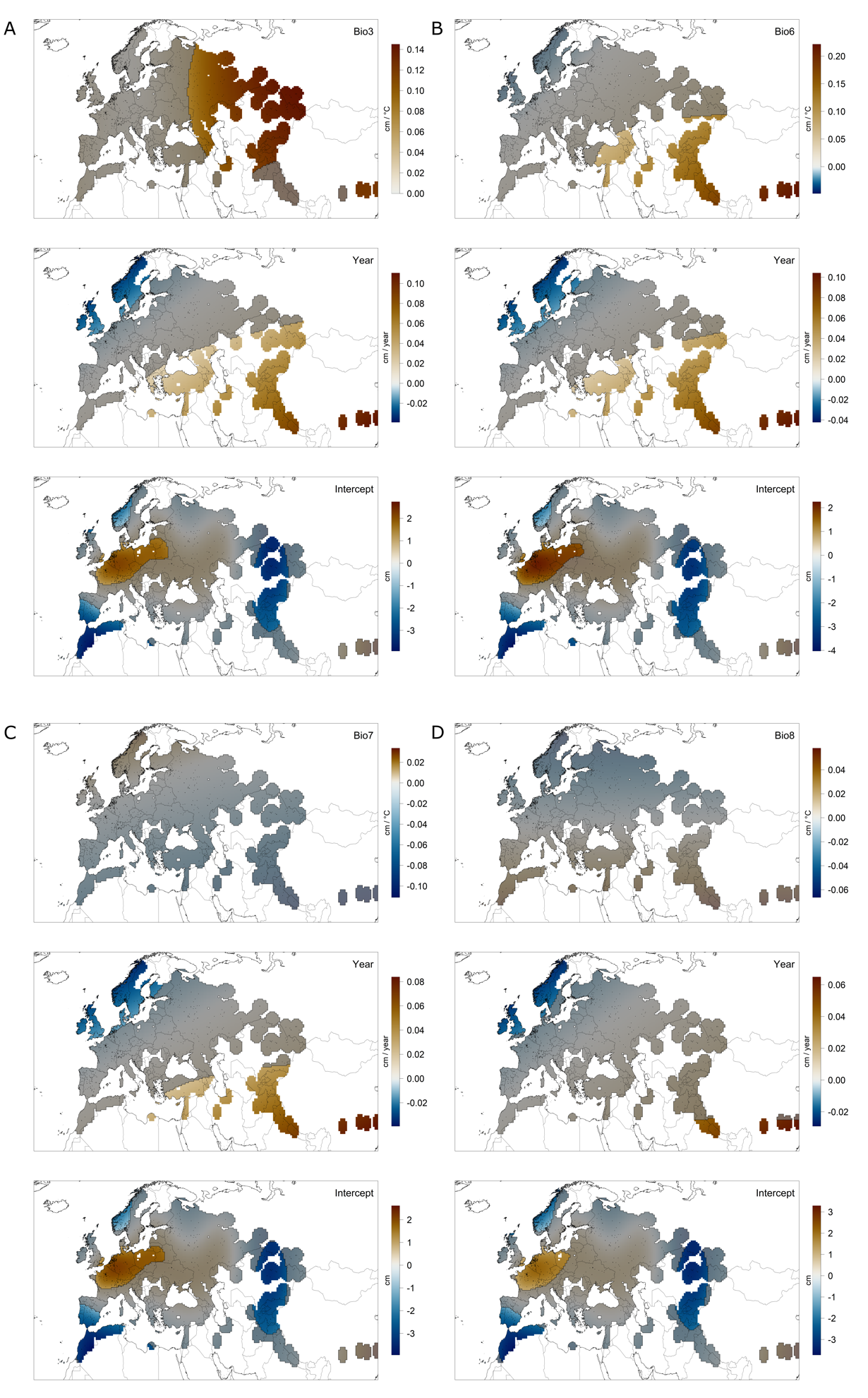

**Figure S3** (cont.). Spatial varying coefficients and intercepts from GAMs showing the relationship between inflorescence height and yearly anomalies in climate variables mean daily mean air temperatures of the warmest quarter (E, BIO10), precipitation seasonality (F, BIO15), mean monthly precipitation amount of the wettest quarter (G, BIO16), and mean monthly precipitation amount of the driest quarter (H, BIO17). Gray shading indicates areas where 95% confidence intervals overlap with 0.

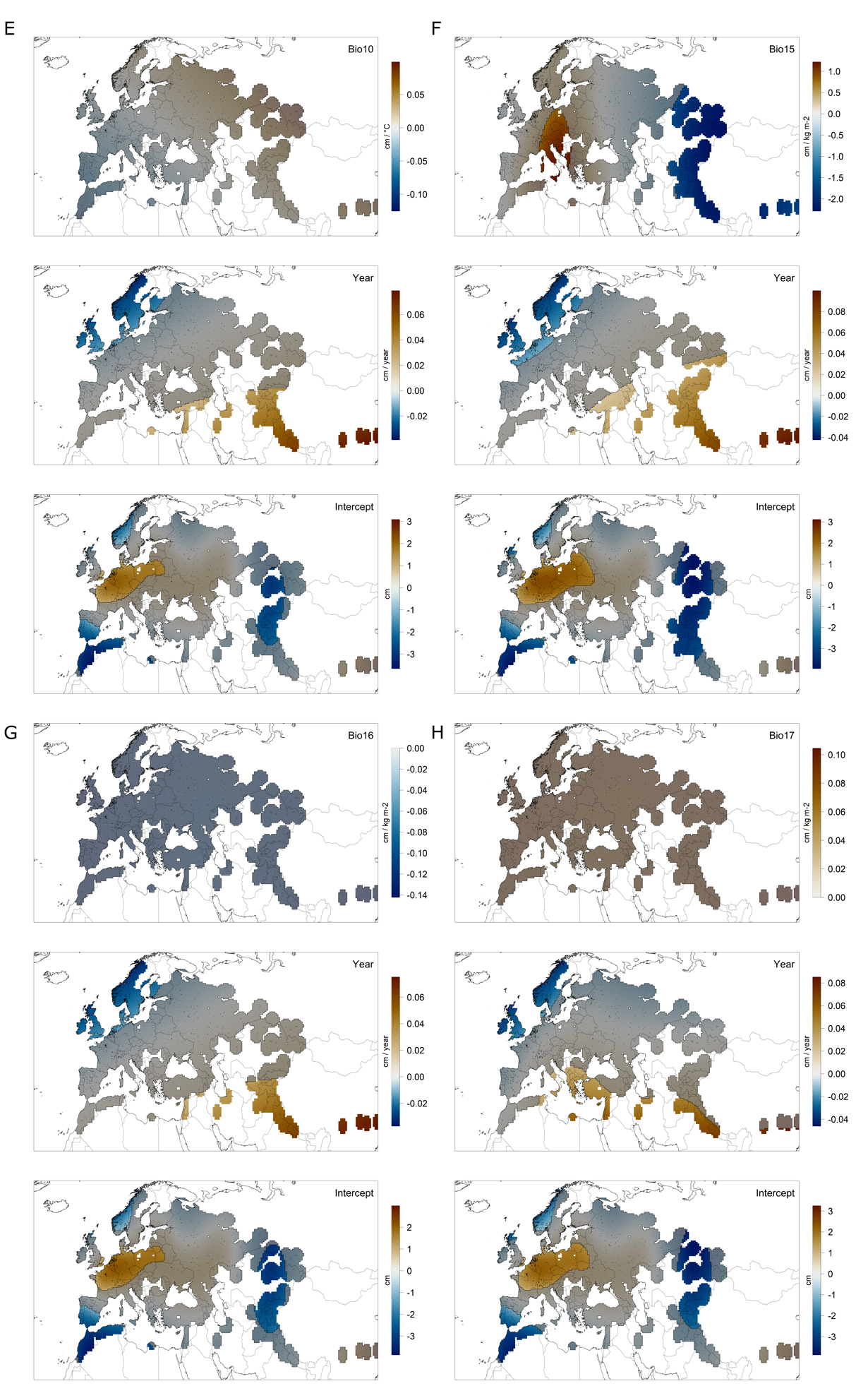

**Figure S4**. Spatial varying coefficients and intercepts from GAMs showing the relationship between maximum rosette leaf length and yearly anomalies in climate variables of isothermality (A, BIO3), mean daily minimum air temperature of the coldest month (B, BIO6), annual range of air temperature (C, BIO7), and mean daily mean air temperature of the wettest quarter (D, BIO8). Gray shading indicates areas where parameter values are not statistically significantly different from zero (e.g. central Europe in for the intercept parameter in panel A), specifically where 95% confidence intervals overlap with 0.

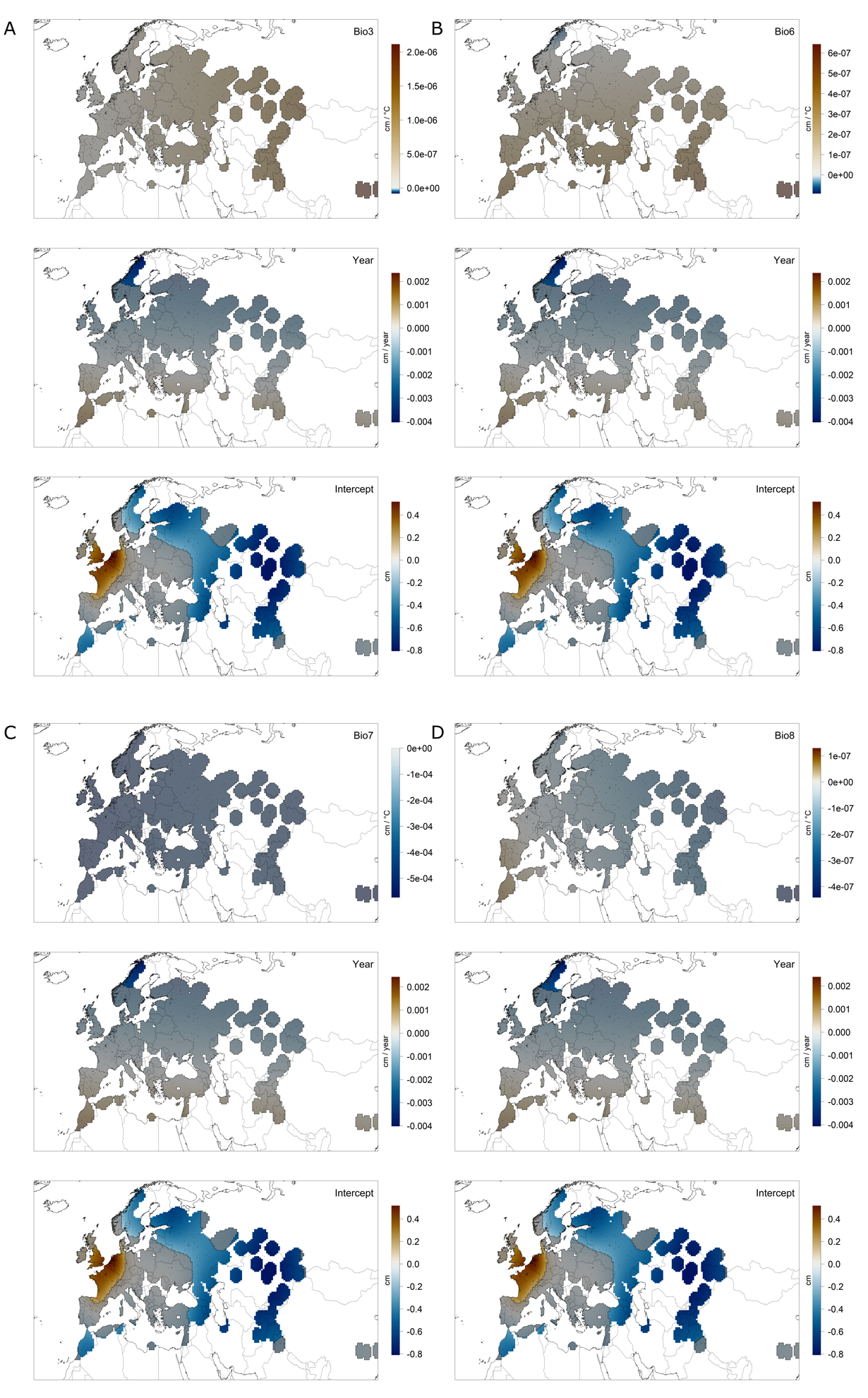
**Figure S4 (cont.).** Spatial varying coefficients and intercepts from GAMs showing the relationship between maximum rosette leaf length and yearly anomalies in climate variables mean daily mean air temperatures of the warmest quarter (E, BIO10), precipitation seasonality (F, BIO15), mean monthly precipitation amount of the wettest quarter (G, BIO16), and mean monthly precipitation amount of the driest quarter (H, BIO17). Gray shading indicates areas where 95% confidence intervals overlap with 0.

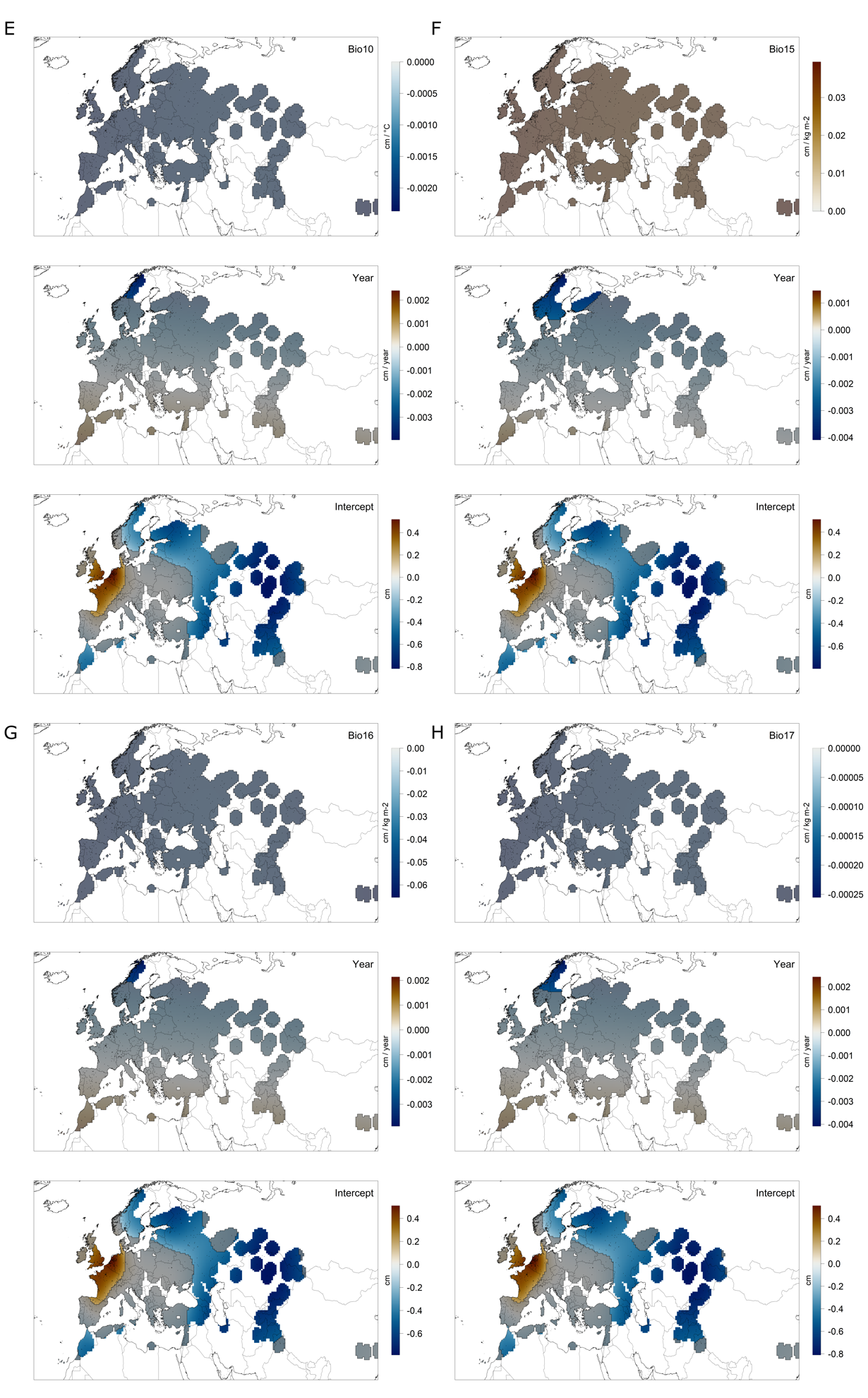

**Figure S5.** The relationship between maximum rosette leaf length and habitat suitability is positive everywhere and significant in western and central Europe in a GAM with spatially varying coefficients. Color indicates fitted parameter values and gray shading covers not statistically significant areas, i.e. where the 95% CI includes zero.

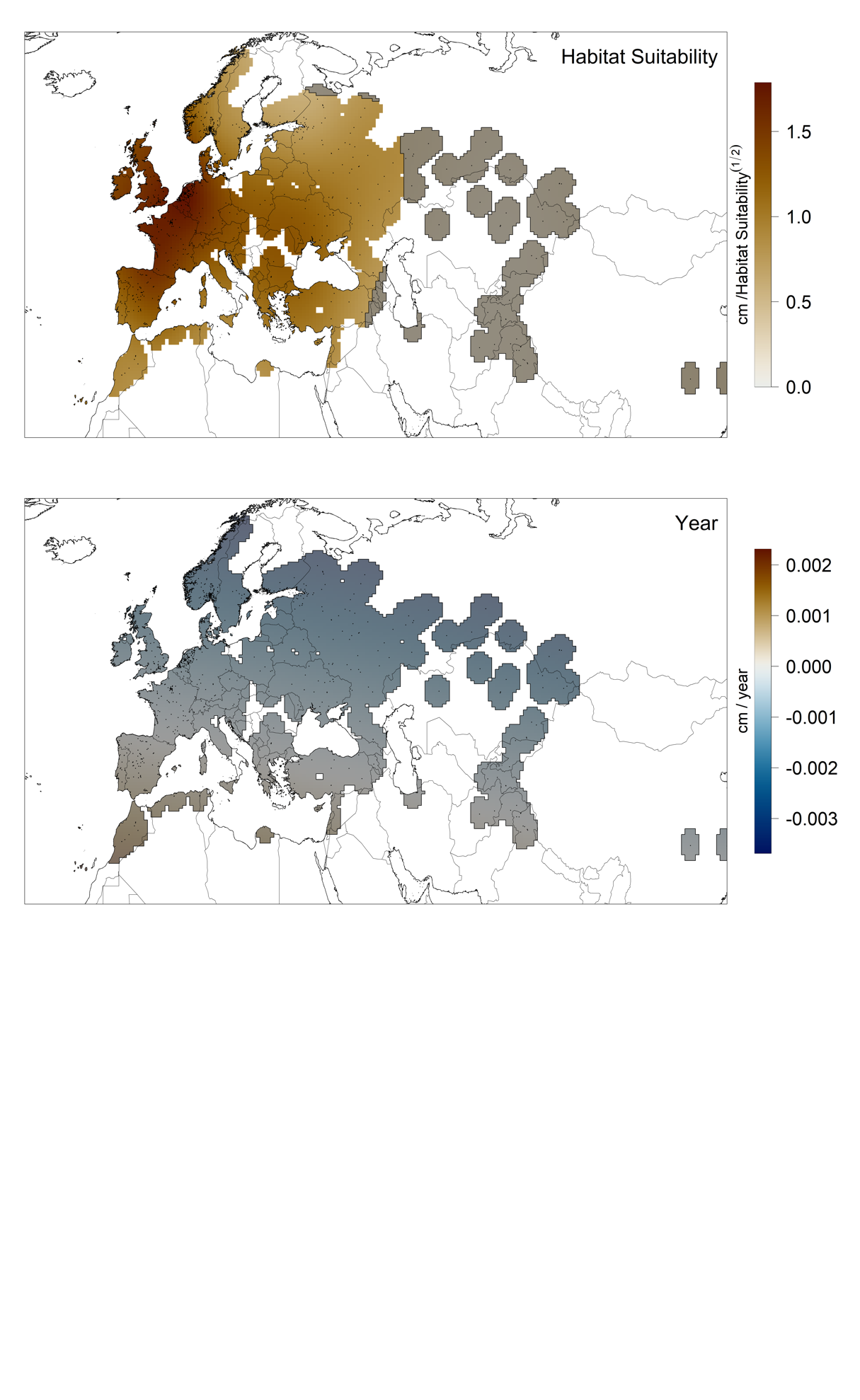

**Figure S6.** The relationship between inflorescence height and habitat suitability is significantly positive across most of Eurasia in a GAM with spatially varying coefficients. Color indicates fitted parameter values and gray shading covers not statistically significant areas, i.e. where the 95% CI includes zero. Additionally, the model indicates plants getting smaller through time in Scandinavia, Britain, and Ireland (blue in bottom panel).

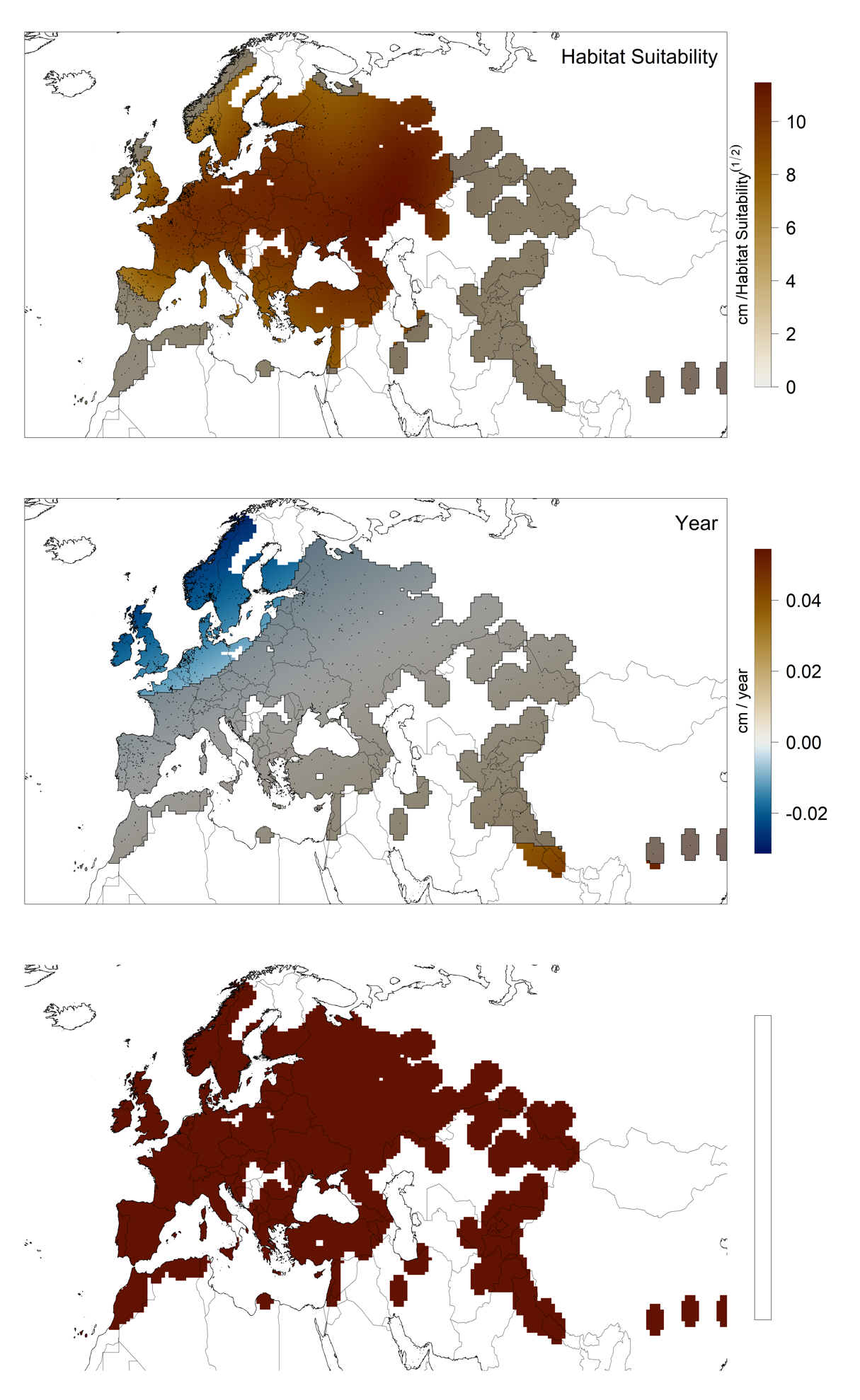

**Figure S7**. Suitability relationship with genetic variation in flowering time (N=953 for days to flower at 10ºC, and N=920 for days to flower at 16ºC and plasticity). Linear regression fits with their associated SE are shown.

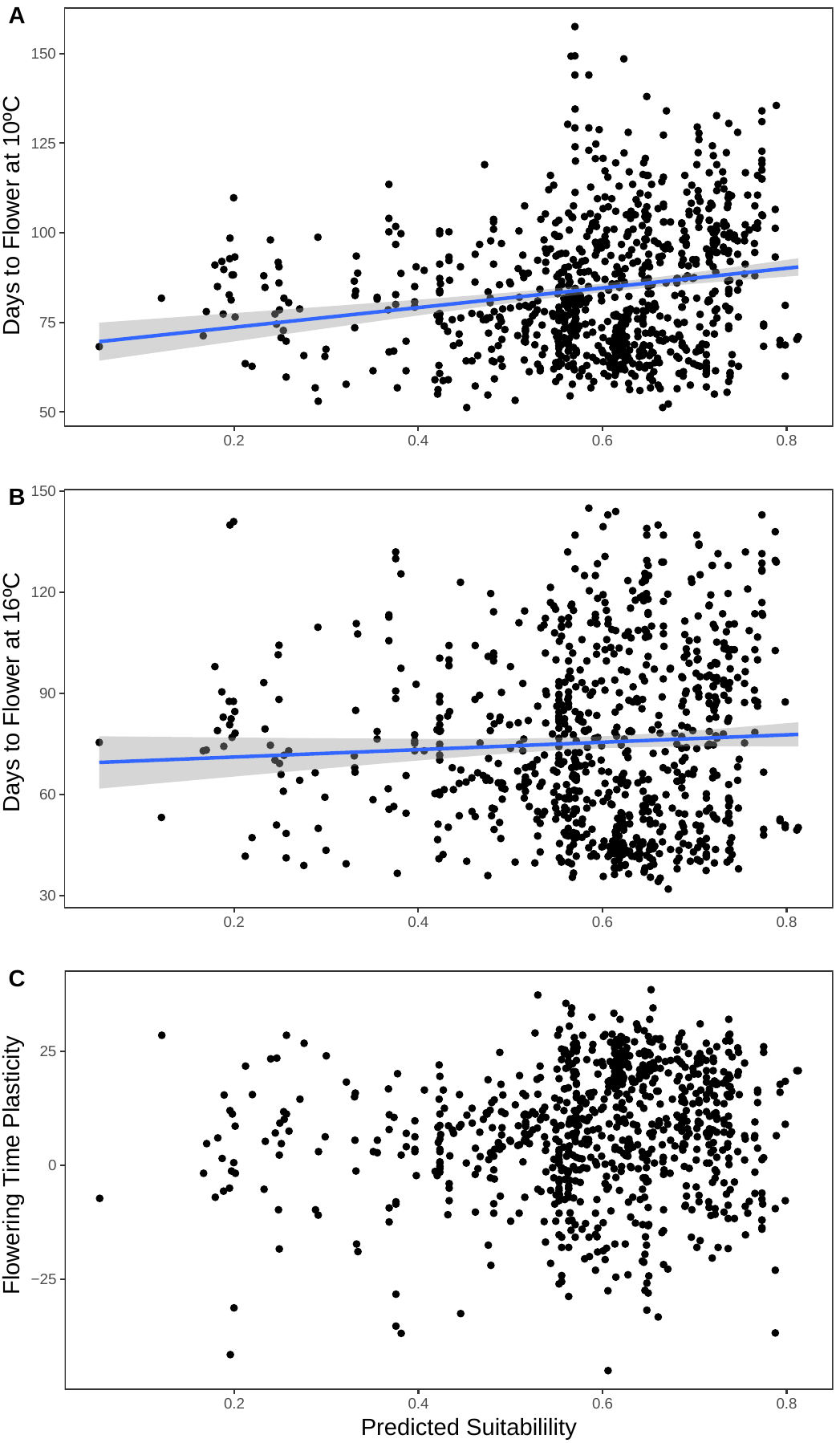

**Figure S8**. Association between habitat suitability and flowering time at 10°C (top) or plasticity of flowering time (bottom) in a GAM. Flowering time was significantly positively related to habitat suitability for most of the locations where Arabidopsis has been collected, while plasticity in flowering time (difference between flowering time at 10°C vs 16°C) is significantly negatively related to habitat suitability. Gray shading indicates regions where parameters are not significantly different from zero, i.e. where the 95% confidence intervals overlap with 0.

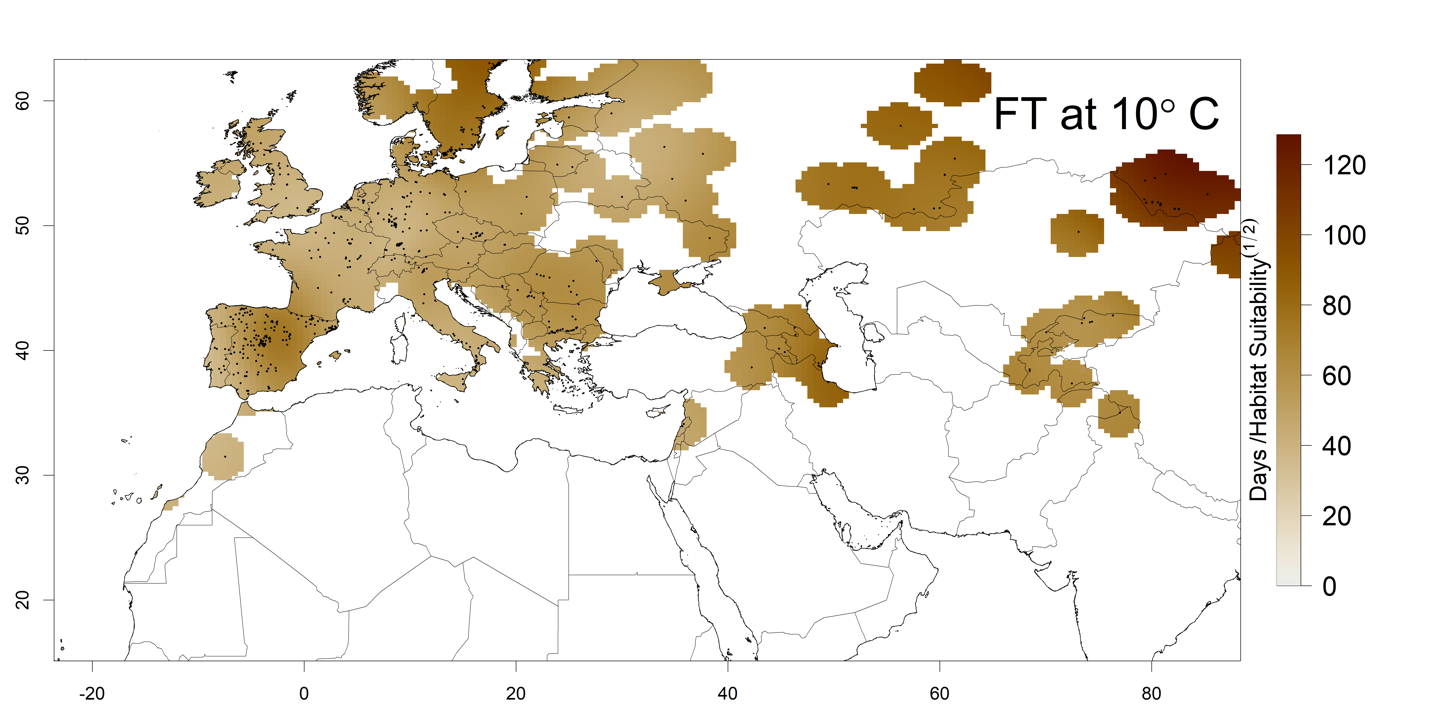

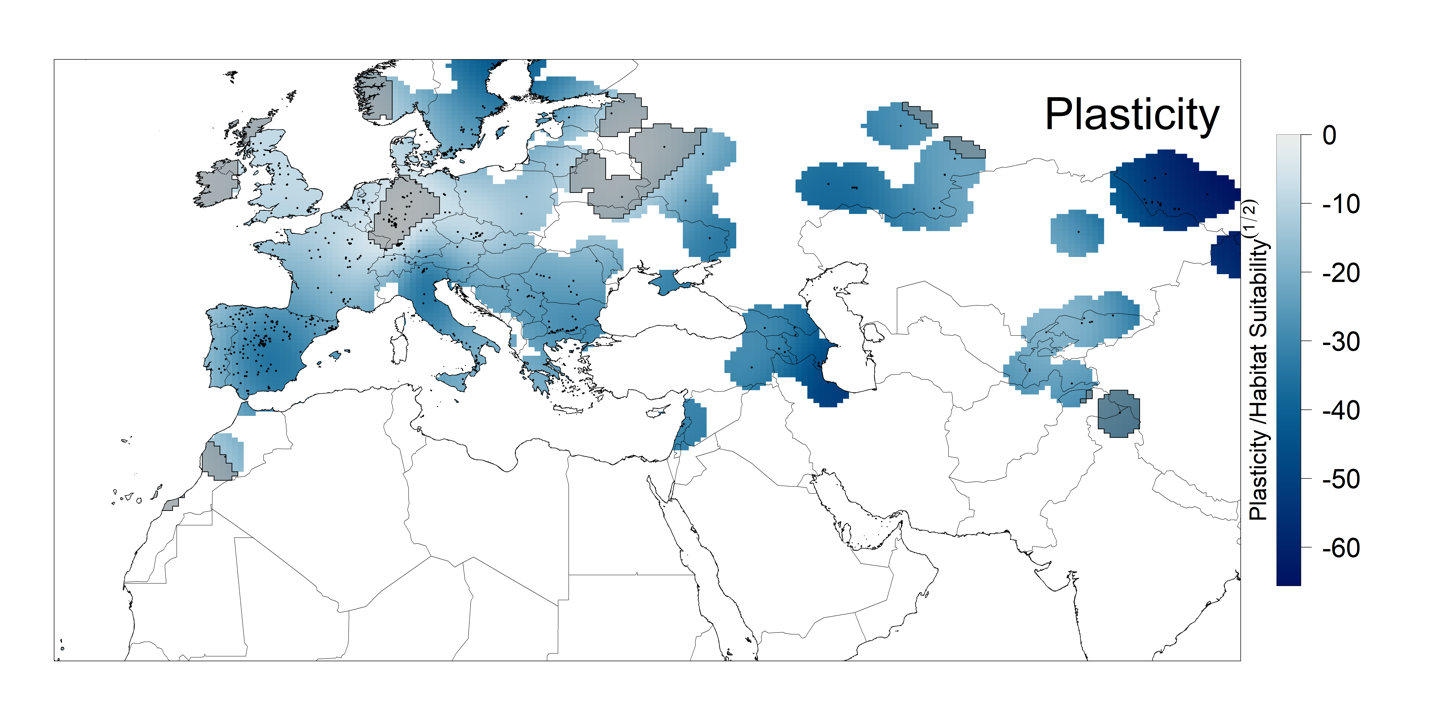

**Figure S9**. Variation in suitability among genetic clusters. Admixed (N=119 ecotypes), Asia (69), Central Europe (168), Germany (54), Italy-Balkan-Caucasus (80), North Sweden (64), South Sweden (153), Relict (24), Spain (110), and Western Europe (92).

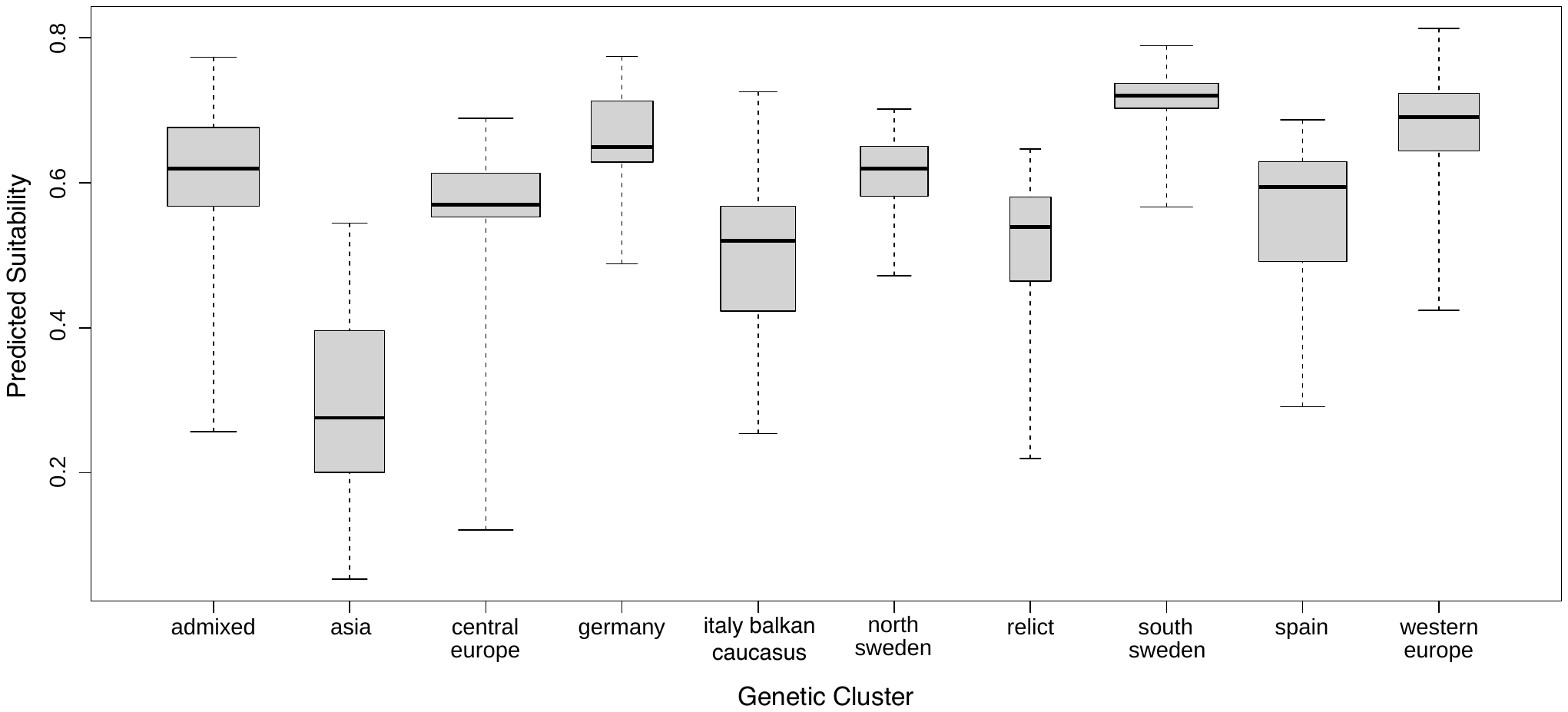

**Figure S10**. Distribution of SNP alleles at the SNP (chr. 2, 8985825 bp) most strongly associated with predicted habitat suitability, found in the putative promoter region (134 bp from the start) of ERF53 (AT2G20880).

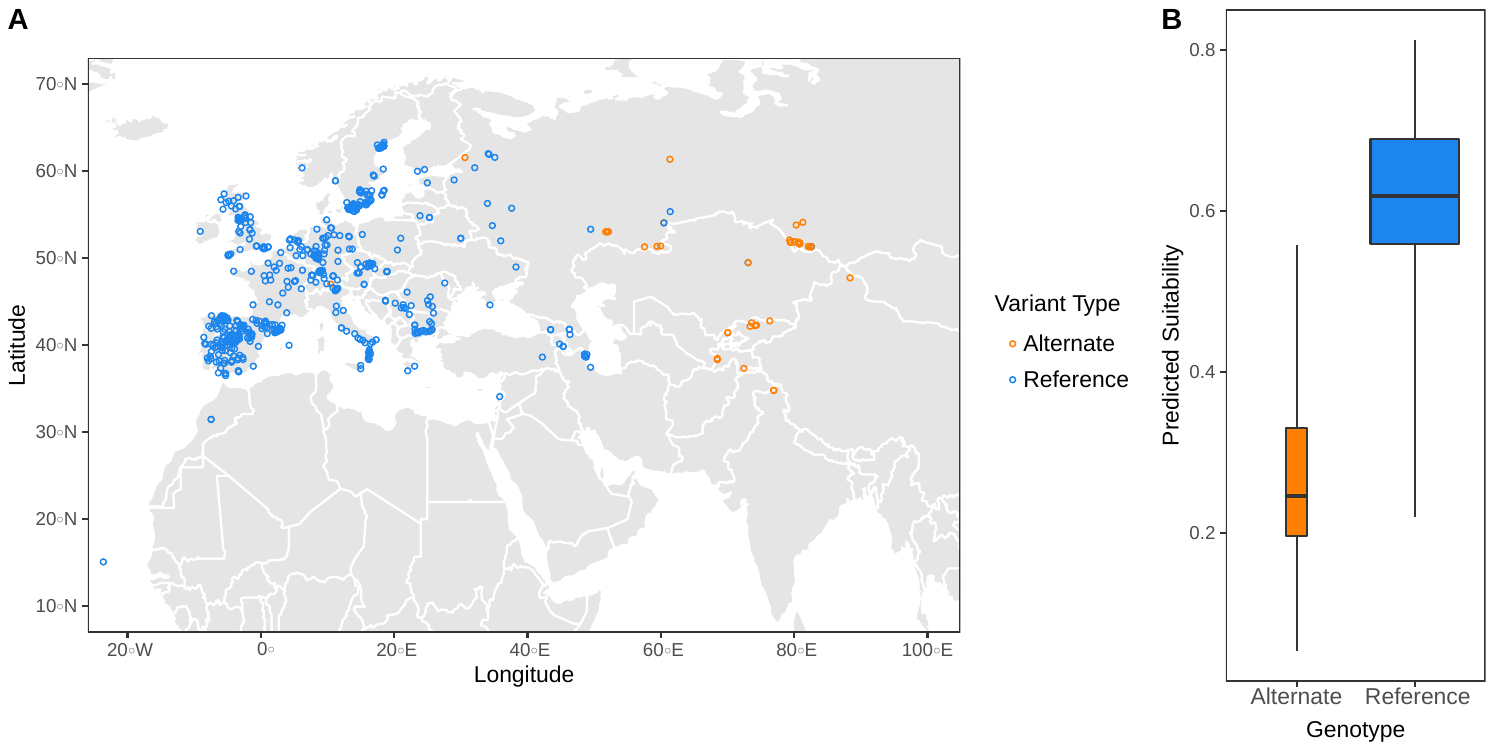

**Figure S11.** Genome-wide associations between Arabidopsis SNPs and habitat suitability, with each of the five chromosomes along the x-axis shown in alternating colors. The top SNP is chr. 2, 8985825 bp, shown in Figure S10. Horizontal dashed line corresponds to a Bonferroni threshold.

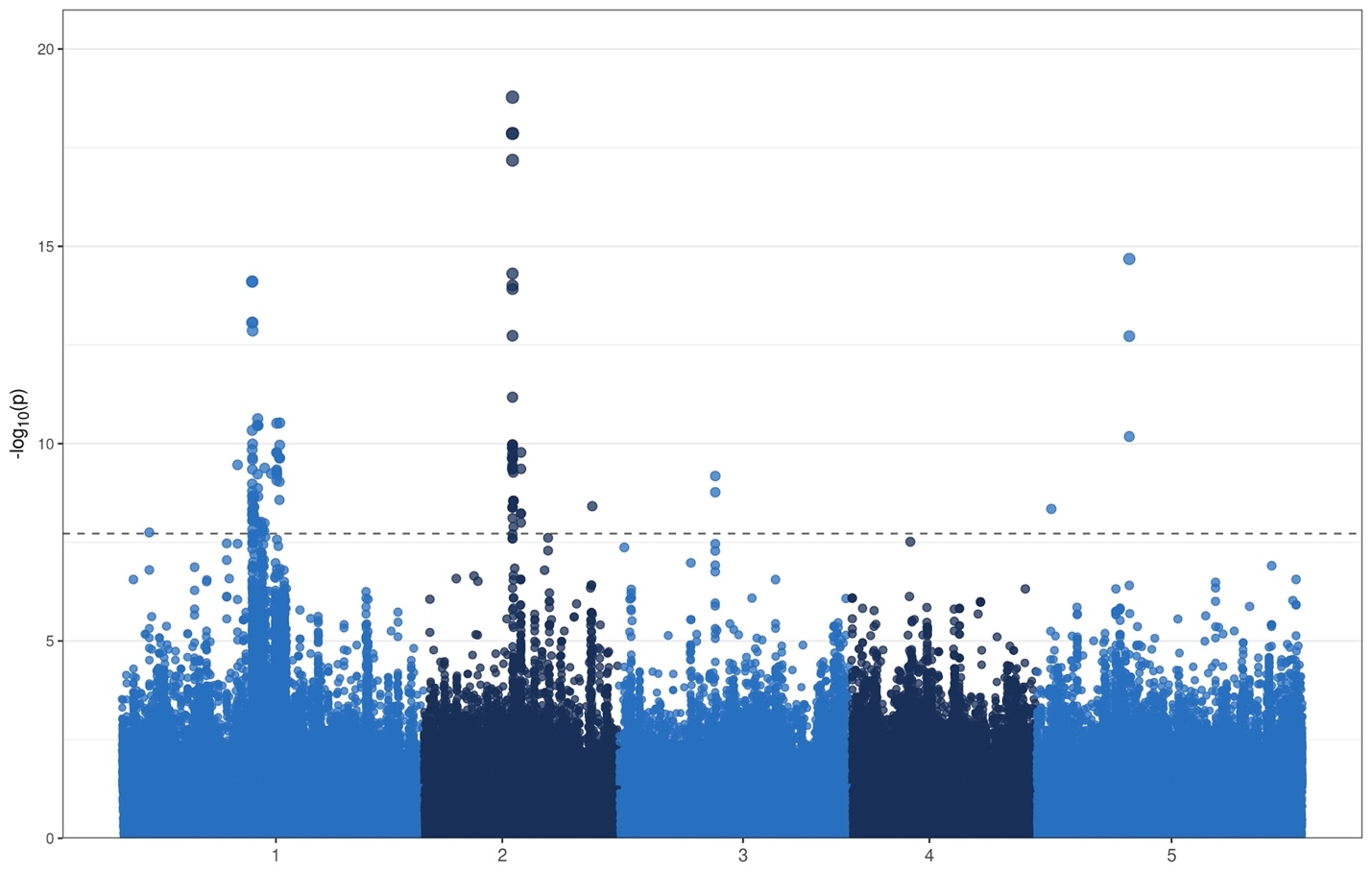

**Figure S12.** Limiting factors map for globe.

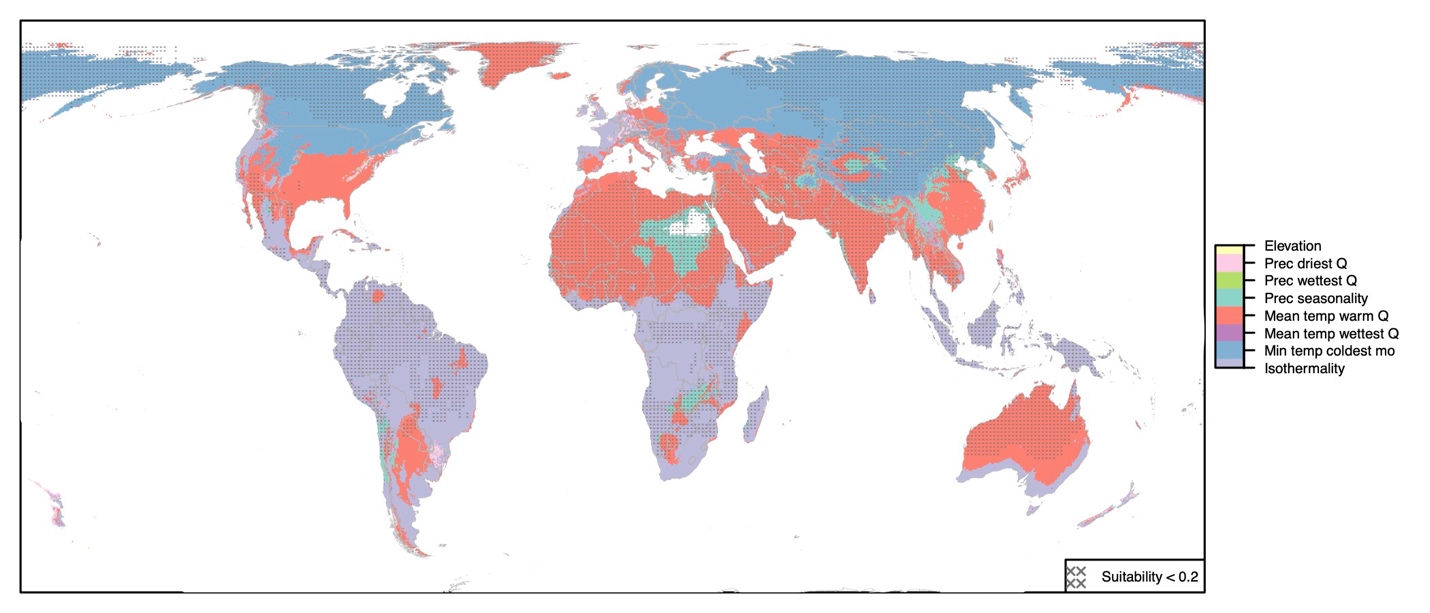

**Figure S13.** Arabidopsis range projections for the future (2050) under RCP 6.0 emissions scenario.

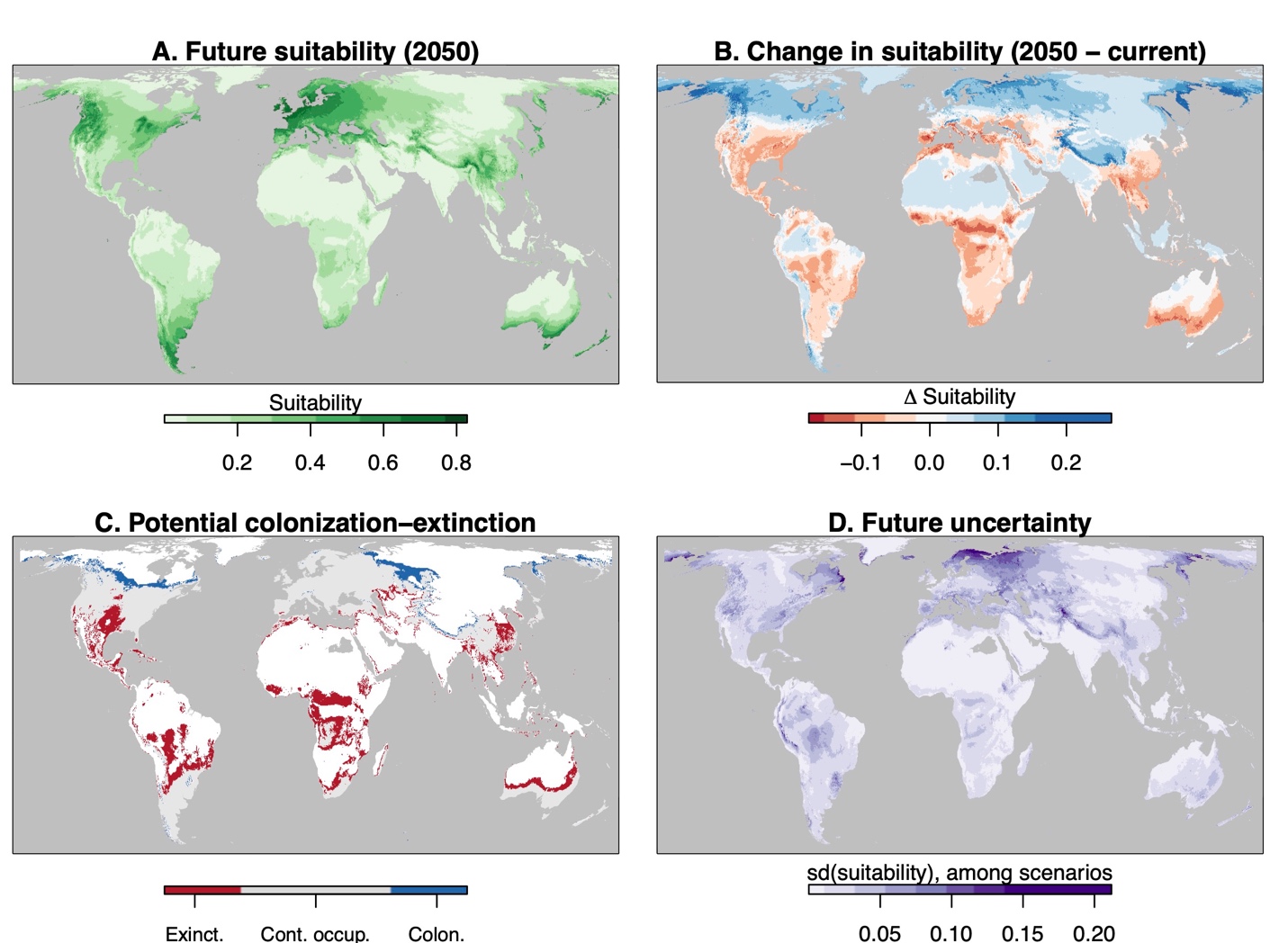

**Figure S14.** (A) Predicted distribution during the Last Glacial Maximum and (B) areas in black with suitability > 0.2 during both the LGM and current conditions as well as within 500 km of known current occurrences. Because of their greater level at the LGM, in (A) we masked the LGM Caspian and Aral Seas (including regions of high putative suitability) from the map (Prentice et al., 1993). Lake Victoria was left unmasked as it was likely very low during the LGM (Johnson et al., 1996). Equal Earth projection was used. For (B), the threshold of 0.2 was chosen for reference as it approximately matches the current distribution of Arabidopsis; the alternate thresholds for visualization of 0.25 and 0.3 and be found in Figures 4 and S15.

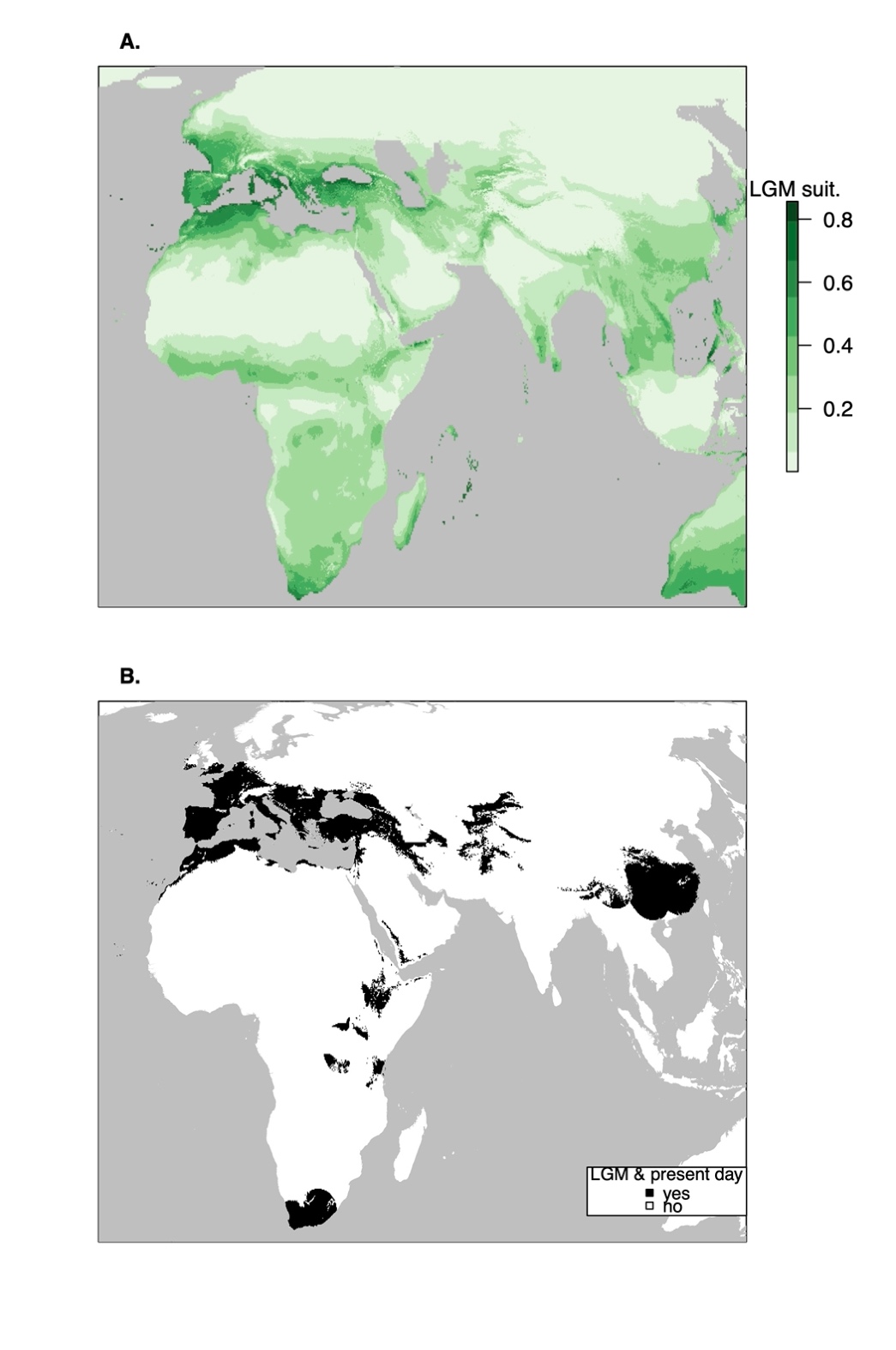

**Figure S15.** (A) Predicted distribution during the Last Glacial Maximum and (B) areas in black with suitability > 0.3 during both the LGM and current conditions as well as within 500 km of known current occurrences. Because of their greater level at the LGM, in (A) we masked the LGM Caspian and Aral Seas (including regions of high putative suitability) from the map (Prentice et al., 1993). Lake Victoria was left unmasked as it was likely very low during the LGM (Johnson et al., 1996). Equal Earth projection was used. For (B), the threshold of 0.3 was chosen for reference as it approximately matches the current distribution of Arabidopsis; the alternate thresholds for visualization of 0.2 and 0.25 and be found in Figures S14 and 4.

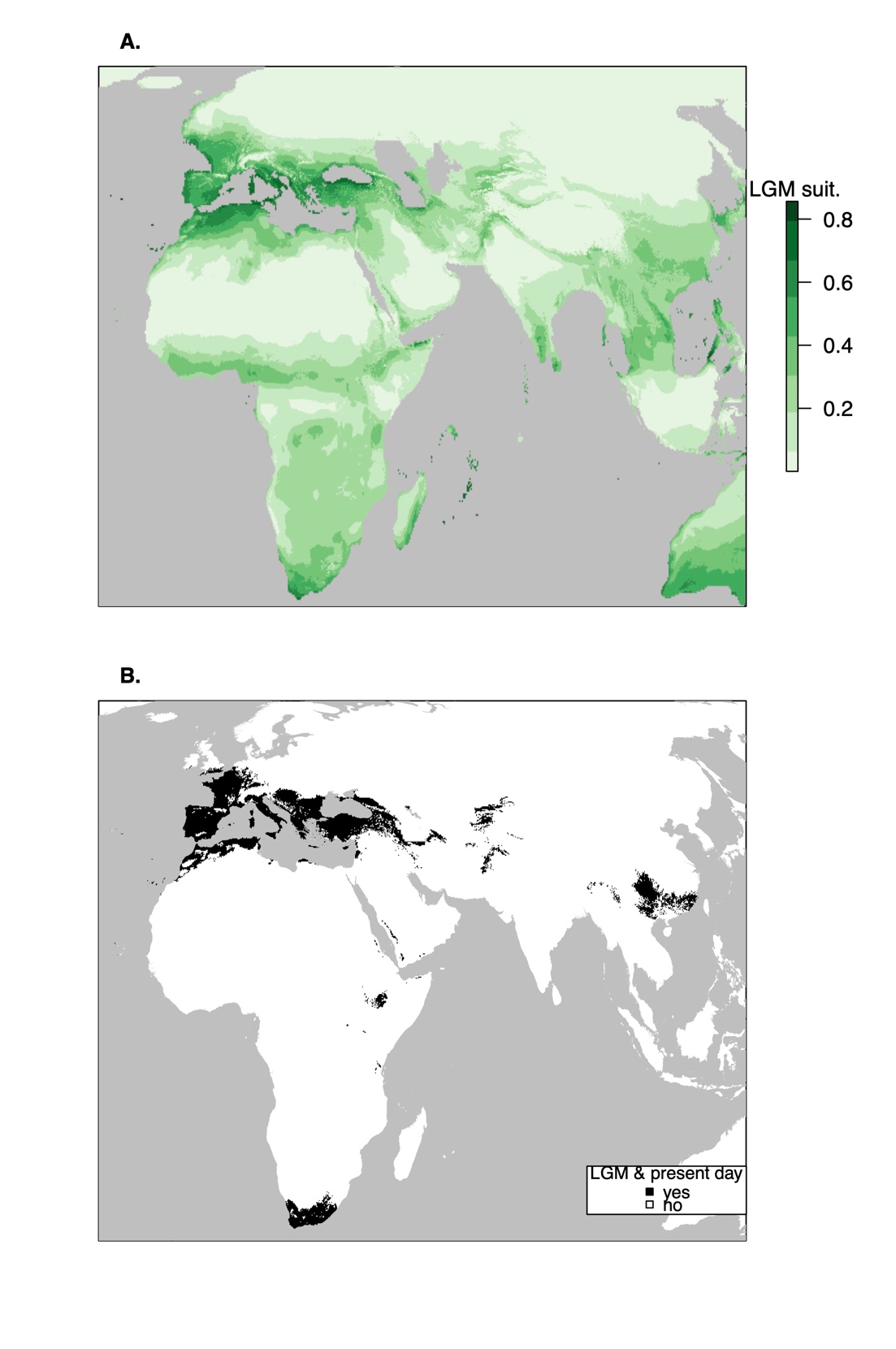

**Figure S16.** Future predicted suitability for Arabidopsis (A), change in suitability from future compared with present (blue indicates improving suitability and red decreasing, B), regions of potential colonization (blue) continued occupancy (gray), and extinction (red) based on a threshold suitability of 0.2 for occupancy (C), and the standard deviation in suitability among the 5 tested climate models giving uncertainty (D). Equal Earth projection is used. RCP 4.5 emissions scenario is shown. For (C), the threshold of 0.2 was chosen for reference as it approximately matches the current distribution of Arabidopsis; the alternate thresholds for visualization of 0.25 and 0.3 and be found in Figures 5 and S17.

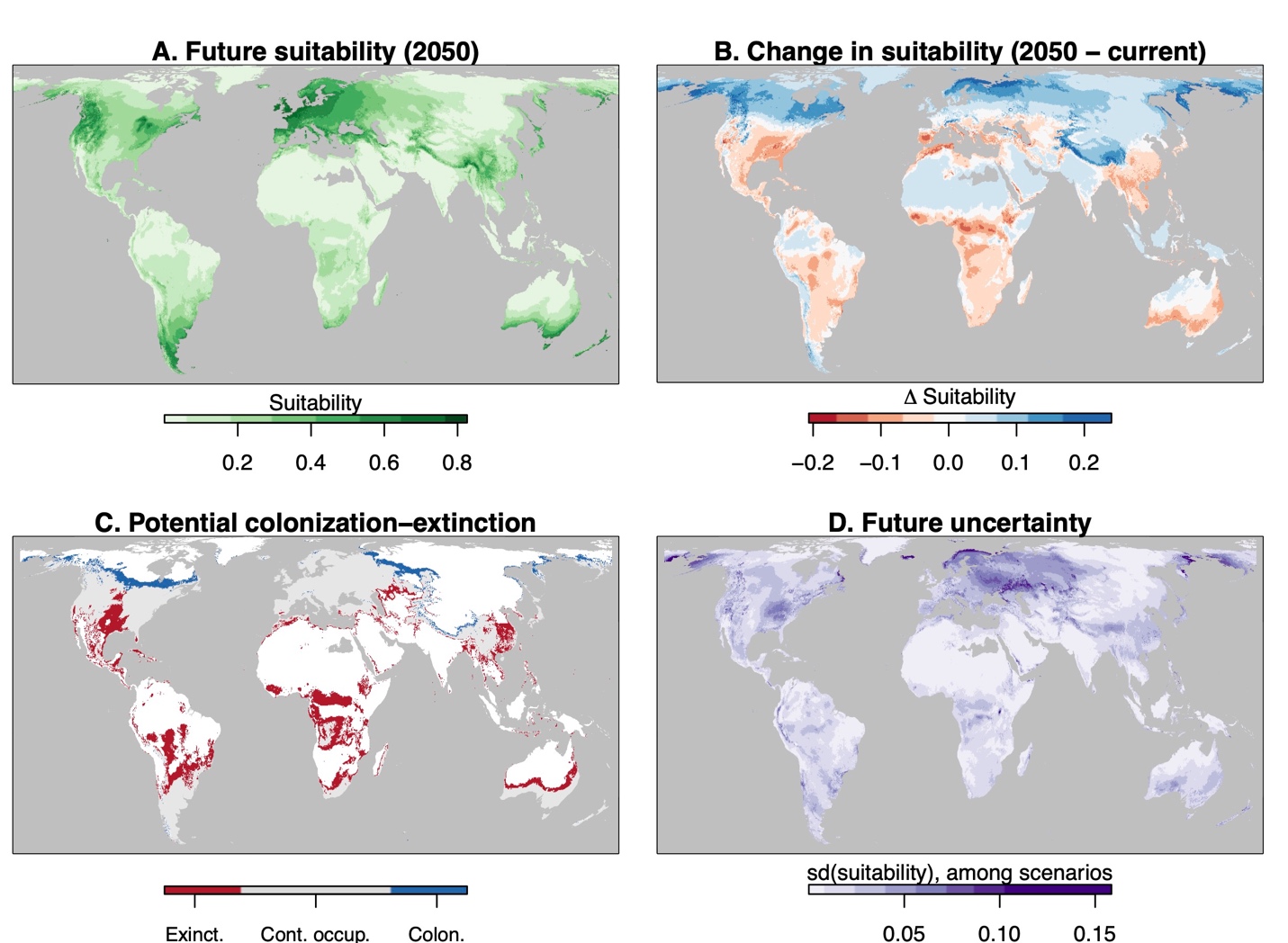

**Figure S17.** Future predicted suitability for Arabidopsis (A), change in suitability from future compared with present (blue indicates improving suitability and red decreasing, B), regions of potential colonization (blue) continued occupancy (gray), and extinction (red) based on a threshold suitability of 0.3 for occupancy (C), and the standard deviation in suitability among the 5 tested climate models giving uncertainty (D). Equal Earth projection is used. RCP 4.5 emissions scenario is shown. For (C), the threshold of 0.3 was chosen for reference as it approximately matches the current distribution of Arabidopsis; the alternate thresholds for visualization of 0.2 and 0.25 and be found in Figures 5 and S16.

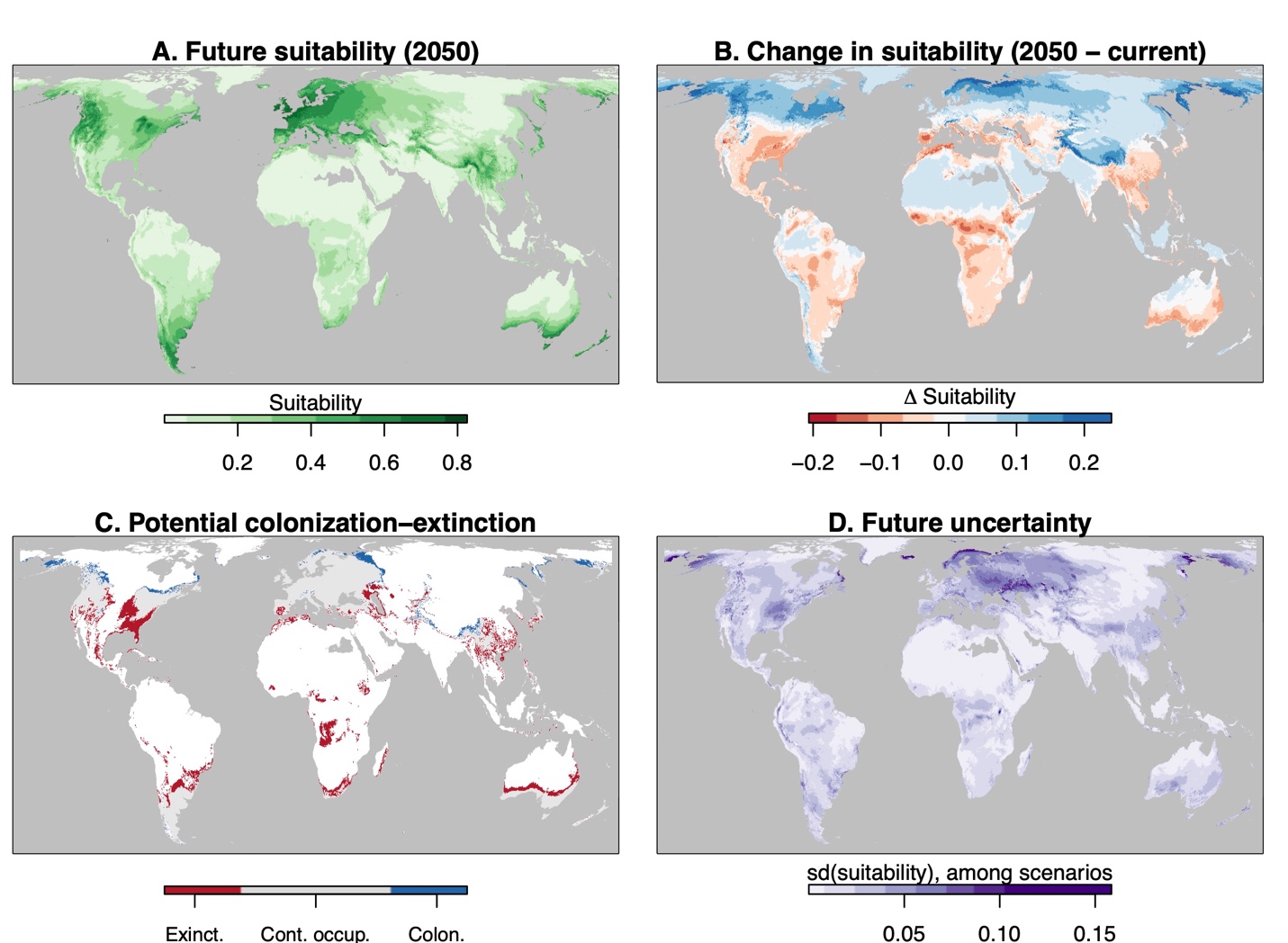

**Figure S18.** Habitat suitability (green) under current climate conditions with thinned occurrences (n = 662) used in fitting shown as black circles. Regions far (>1000 km) from known occurrence have a gray mask. Compare with Figure 1A which shows the same output except for a model fit using a 500 km buffer for background pseudoabsences. Equal Earth projection was used.

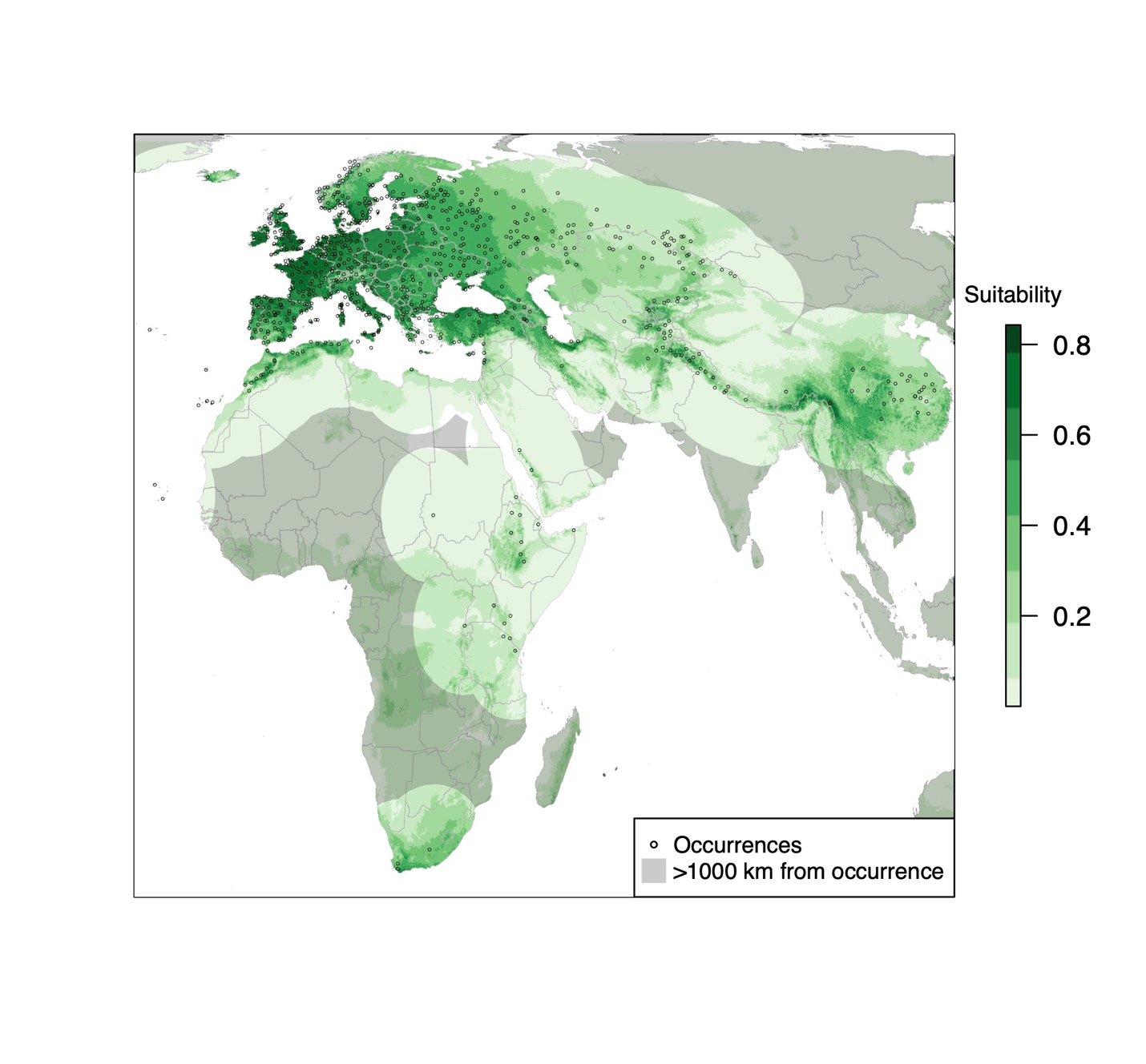

**Table S1**. Permutation importance of tested variables in the fitted habitat suitability model.

| Environmental Variable | Permutation Importance (%) |
| --- | --- |
| Min Temperature of Coldest Month | 38.1515 |
| Mean Temperature of Warmest Quarter | 31.1246 |
| Isothermality | 15.2931 |
| Precipitation Seasonality | 9.5232 |
| Elevation | 3.3047 |
| Mean Temperature of Wettest Quarter | 1.6697 |
| Precipitation of Driest Quarter | 0.7389 |
| Temperature Annual Range | 0.1879 |
| Precipitation of Wettest Quarter | 0.0066 |

**Table S2**. Results from linear mixed-effects regressions fit by maximum likelihood of life history variables (days to flower at 10ºC and 16ºC, and flowering time plasticity) on predicted suitability. Models account for kinship among ecotypes (random effect) using the R package coxme (function lmekin). Significant relationships (*P* ≤ 0.05) are bolded.

|  |  | Response | | |
| --- | --- | --- | --- | --- |
|  |  | DTF^a^ 10ºC | DTF 16 ºC | plasticity |
|  | N | 884 | 852 | 852 |
|  | log-likelihood | -3408.44 | -3580.2 | -3230.61 |
| Intercept | coefficient | 76.75 | 71.09 | 5.89 |
|  | Std Error | 21.65 | 30.46 | 20.21 |
|  | *Z* | 3.54 | 2.33 | 0.29 |
|  | *P* | 0.0004 | 0.02 | 0.77 |
| Fixed  Effect | coefficient | 19.09 | 19.58 | -1.52 |
|  | Std Error | 6.06 | 8.6 | 5.71 |
|  | *Z* | 3.15 | 2.28 | -0.27 |
|  | *P* | **0.002** | **0.02** | 0.79 |
| Random  Effect | Std Dev | 4.00E+08 | 1.59E+09 | 23.54 |
|  | variance | 1.57E+17 | 2.54E+18 | 554.12 |

^a^DTF: days to flower

**Table S5**. Tukey HSD post-hoc tests for genetic clusters’ predicted suitability. Genetic clusters are: Admixed (N=119 ecotypes), Asia (69), Central Europe (168), Germany (54), Italy-Balkan-Caucasus (80), North Sweden (64), South Sweden (153), Relict (24), Spain (110), and Western Europe (92). Significant comparisons (*P* ≤ 0.05) are bolded.

|  | difference | lower | upper | P adjusted |
| --- | --- | --- | --- | --- |
| Asia - Admixed | -0.30 | -0.34 | -0.26 | **5.118E-13** |
| Central Europe - Admixed | -0.04 | -0.07 | -0.01 | **2.355E-04** |
| Germany - Admixed | 0.05 | 0.01 | 0.09 | **7.941E-04** |
| Italy Balkan Caucasus - Admixed | -0.10 | -0.14 | -0.07 | **6.052E-13** |
| North Sweden - Admixed | 0.00 | -0.03 | 0.04 | 1.000E+00 |
| Relict - Admixed | -0.10 | -0.15 | -0.04 | **5.354E-07** |
| South Sweden - Admixed | 0.10 | 0.07 | 0.13 | **6.043E-13** |
| Spain - Admixed | -0.05 | -0.08 | -0.02 | **3.262E-05** |
| Western Europe - Admixed | 0.07 | 0.04 | 0.11 | **9.993E-10** |
| Central Europe - Asia | 0.26 | 0.22 | 0.29 | **5.118E-13** |
| Germany - Asia | 0.35 | 0.31 | 0.40 | **5.118E-13** |
| Italy Balkan Caucasus - Asia | 0.19 | 0.16 | 0.23 | **5.118E-13** |
| North Sweden - Asia | 0.30 | 0.26 | 0.35 | **5.118E-13** |
| Relict - Asia | 0.20 | 0.14 | 0.26 | **5.531E-13** |
| South Sweden - Asia | 0.40 | 0.36 | 0.43 | **5.118E-13** |
| Spain - Asia | 0.25 | 0.21 | 0.29 | **5.118E-13** |
| Western Europe - Asia | 0.37 | 0.33 | 0.41 | **5.118E-13** |
| Germany - Central Europe | 0.10 | 0.06 | 0.13 | **7.804E-13** |
| Italy Balkan Caucasus - Central Europe | -0.06 | -0.10 | -0.03 | **7.203E-08** |
| North Sweden - Central Europe | 0.05 | 0.01 | 0.08 | **2.428E-03** |
| Relict - Central Europe | -0.06 | -0.11 | 0.00 | **2.537E-02** |
| South Sweden - Central Europe | 0.14 | 0.11 | 0.17 | **5.118E-13** |
| Spain - Central Europe | -0.01 | -0.04 | 0.02 | 9.962E-01 |
| Western Europe - Central Europe | 0.11 | 0.08 | 0.14 | **5.226E-13** |
| Italy Balkan Caucasus - Germany | -0.16 | -0.20 | -0.12 | **5.129E-13** |
| North Sweden - Germany | -0.05 | -0.10 | -0.01 | **1.338E-02** |
| Relict - Germany | -0.15 | -0.21 | -0.09 | **6.973E-13** |
| South Sweden - Germany | 0.05 | 0.01 | 0.08 | **6.121E-03** |
| Spain - Germany | -0.10 | -0.14 | -0.06 | **6.598E-13** |
| Western Europe - Germany | 0.02 | -0.02 | 0.06 | 9.355E-01 |
| North Sweden - Italy Balkan Caucasus | 0.11 | 0.07 | 0.15 | **6.143E-13** |
| Relict - Italy Balkan Caucasus | 0.01 | -0.05 | 0.06 | 1.000E+00 |
| South Sweden - Italy Balkan Caucasus | 0.20 | 0.17 | 0.24 | **5.118E-13** |
| Spain - Italy Balkan Caucasus | 0.05 | 0.02 | 0.09 | **5.561E-05** |
| Western Europe - Italy Balkan Caucasus | 0.18 | 0.14 | 0.21 | **5.118E-13** |
| Relict - North Sweden | -0.10 | -0.16 | -0.04 | **1.554E-06** |
| South Sweden - North Sweden | 0.10 | 0.06 | 0.13 | **6.133E-13** |
| Spain - North Sweden | -0.05 | -0.09 | -0.02 | **3.656E-04** |
| Western Europe - North Sweden | 0.07 | 0.03 | 0.11 | **2.120E-06** |
| South Sweden - Relict | 0.20 | 0.14 | 0.25 | **5.129E-13** |
| Spain - Relict | 0.05 | -0.01 | 0.10 | 1.423E-01 |
| Western Europe - Relict | 0.17 | 0.11 | 0.23 | **6.076E-13** |
| Spain - South Sweden | -0.15 | -0.18 | -0.12 | **5.118E-13** |
| Western Europe - South Sweden | -0.03 | -0.06 | 0.00 | 1.492E-01 |
| Western Europe - Spain | 0.12 | 0.09 | 0.16 | **5.332E-13** |
